# Foldseek-Interface reveals a protein interface universe far from complete

**DOI:** 10.64898/2026.08.24.746585

**Authors:** Joelle Morgan Strom, Sooyoung Cha, Rachel Seongeun Kim, Harshit Sajal, Cameron L. M. Gilchrist, Martin Steinegger, Katja Luck

## Abstract

Protein-protein interactions mediate a vast range of cellular functions, requiring diverse modes of binding. While recent years have seen major efforts to chart and classify the protein structure universe, we lack comparable methods to assess and cluster that diversity in interface structure at interactome scale. Here, we present Foldseek-Interface, a method that converts 3D interface structures into searchable sequences to enable fast alignment and clustering of protein interaction interfaces. It matches the accuracy of state-of-the-art tools while running up to 230 times faster. Applying it to all biological assemblies in the PDB, we cluster 3.1 million dimers into 77,167 interface clusters and use this resource to characterise interface diversity, evolution, and pathogen mimicry. Application of Foldseek-Interface to resources of predicted protein complex structures rapidly revealed putatively novel interface types worth further experimental interrogation. Foldseek-Interface and the interface cluster resource are freely available as webservers for search (https://search.foldseek.com/interface) and exploration (https://interface.foldseek.com).

---

Protein-protein interactions (PPIs) drive cellular life by mediating responses to extra- and intracellular stimuli, orchestrating development and enabling organismal adaptation. This functional diversity depends on highly dynamic modes of protein association and dissociation that remain incompletely understood. Proteins contain regions that independently fold into tertiary structures (ordered regions) (1) and regions that remain unstructured in their unbound state, termed intrinsically disordered regions (IDRs) (2). Interactions between ordered protein regions and coiled coils mediate stable complexes and rigid scaffolds, whereas interactions involving disordered regions, often through short linear motifs that bind ordered regions in partner proteins, form transient signaling complexes and mediate condensation (3). The 3D coordinates of residues in both interacting protein chains that are in contact with each other as well as other close-by residues form an interface structure.

Recent advances in accurate protein structure prediction (4) and fast structure search (5) have enabled systematic analysis of the protein fold space, revealing 2.3 million structural clusters (6). However, equivalent efforts to chart the resolved and predicted universe of interface structures are missing. Similar interface structures can occur across otherwise unrelated complexes, through convergent evolution (7). Therefore, comparing interfaces directly, rather than entire complexes, is needed to recognize shared binding modes, and clustering interfaces by structural similarity would organize the known interactome into recurring topological families (8, 9). Doing this across the Protein Data Bank (PDB) (10) requires a method that can both align and cluster interfaces efficiently regardless of interface size and type. Previous studies were often restricted to larger, ordered interfaces, combined overall sequence and complex similarity with interface similarity searches, or performed geometric motif searches(11–14). iAlign (15) is a state-of-the-art tool for computing local interface similarity that is amenable to ordered and disordered interfaces, yet, would take approximately 50 years to compute pairwise similarity across all interface structures in the PDB. This challenge will only increase as thousands of predicted protein complex structures are already available (16–19), with many more expected in the near future. How can we efficiently search these resources to discover novel predicted modes of protein binding?

The speed bottleneck can be bypassed by reducing 3D structures to 1D strings, thereby turning structural comparison into fast sequence alignments. Foldseek (5) does this by translating each residue into a 3Di structural alphabet state that encodes its structural relationship with the nearest neighbour in space, and aligning the resulting strings. Foldseek-Multimer (20) accelerates chain-to-chain alignments for multi-subunit protein complexes but does not capture local interface similarity. Foldseek’s encoding and alignment assume a single, contiguous chain. An interface, however, is formed between two chains and the interface residues within a chain are not contiguous. Thus, when an interface residue’s nearest neighbour lies on the partner chain, the sequence offset encoded by 3Di loses its meaning across chains, and there is no single N-to-C sequence for order-dependent alignment.

Here, we present Foldseek-Interface, a method that aligns interfaces at high speed and accuracy. It resolves the contiguity problem by renumbering extracted interface residues in N-to-C order within each chain, generating a contiguous 3Di string that Foldseek aligns directly, while Foldseek-Multimer recovers the matching chains. We further develop a clustering framework calibrated on interface-only TM-score (21) thresholds for clustering of interfaces at database-scale. Applying this to all biological assemblies in the PDB, we cluster 3.1 million protein dimers into 77,167 non-redundant interface clusters representing putative interface types and show that interface discovery continues even as fold discovery plateaus. Using this resource, we identify a strong bias in the PDB toward ordered interfaces and explore interface evolution, including convergent evolution and pathogen-host mimicry. We further search predicted human protein complex structures (18) to identify putatively novel interface types and thus, chart, classify, and systematically expand the protein interface space.

## Results

The core innovation of Foldseek-Interface is a 3Di encoding of interface residues rather than of full chains (Fig. 1a-b). Interface residues are identified between interacting chains using an *α*-carbon distance of *≤* 10Å, and chain pairs with *≥* 4 interface residues per chain are retained. Interface residues are then extracted from each chain in N- to C-terminal order, and both chains are concatenated before 3Di encoding. This allows each residue’s nearest neighbor to lie on either the same or the partner chain, thereby encoding the interface geometry defined by both binding partners (Fig. 1b).

**Fig. 1.**
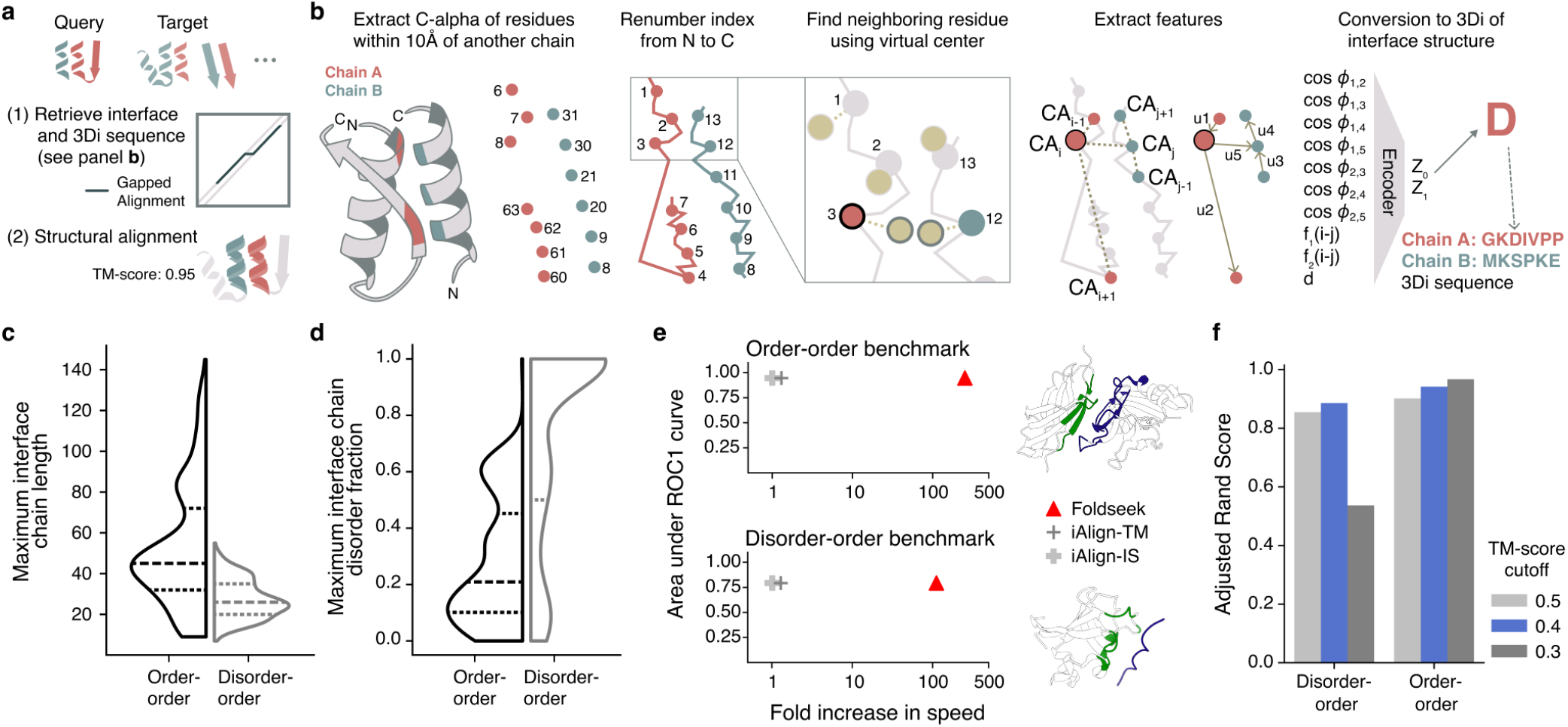
Development and benchmarking of Foldseek-Interface. **(a)** Schematic showing structural alignment of interfaces with Foldseek-Interface. **(b)** Schematic showing the extraction of interface residues and conversion to 3Di sequence. **(c**,**d)** Violin plots comparing the maximum interface chain length (c) and maximum interface chain disorder fraction (d) for the order-order and disorder-order benchmark datasets, Mann-Whitney U-test statistic=372679, 52350; p-value=1.30 *\** 10^−76^ and 2.13 *\** 10^−78^ for interface length and disorder, respectively. **(e)** Scatterplot comparing accuracy of all-by-all interface alignment and scoring vs. run-time for Foldseek-Interface and both scoring methods of iAlign for two benchmark datasets. Speed represented as the fold-increase in speed over the slowest method. Example of order-order structure (1TDQ) and disorder-order structure (3I5R) shown as insets with interfaces highlighted in color. **(f)** Barplot representing the accuracy of Foldseek-Interface-Cluster of the benchmark datasets when clustered within all resolved interfaces from the PDB at various interface TM-score cut-offs.

To benchmark the speed and accuracy of Foldseek-Interface across different modes of protein binding, we created separate benchmark datasets of clustered order-order and disorder-order interfaces (Table S1, S2). Because no geometry-based reference clustering of protein interfaces is available, we used resources that define interface similarity by combining fold and protein sequence similarity (22, 23). The two datasets differ substantially in interface size and disorder content (Fig. 1c-d), providing complementary benchmarks for distinct modes of protein binding. Because interfaces are substantially shorter than full-length structures, we replaced the pre-filtering step with exhaustive alignment and optimized parameters for interface comparison (Fig. S1a). Poorly aligned chain pairs were rejected using an lDDT cutoff of 0.2 before interface alignment and calculation of the interface TM-score (21) (Fig. S1b-c). In all-by-all comparisons, Foldseek-Interface achieved comparable accuracy to iAlign (15) (0.966 vs. 0.949 for order-order; 0.789 vs. 0.809 for disorder-order), while running up to 230 times faster (Fig. 1e, Fig. S1d-e, Table S3, S4). Both methods performed worse on disorder-order interfaces, likely because of their smaller size and higher flexibility.

To enable efficient clustering of protein complexes, we extended a clustering algorithm with four complementary measures of structural similarity, and combined it with Foldseek-Interface to create Foldseek-Interface-Cluster (Fig. S1f, see Methods). Using the order-order and disorder-order benchmark datasets, we evaluated clustering accuracy of both benchmark datasets within the full PDB (see Methods and below) across interface TM-score cutoffs (Fig. 1e, Table S5, S6). This yielded an optimal cutoff of 0.4 (Fig. 1f). Overall, these benchmarks establish Foldseek-Interface as an accurate and scalable method for large-scale structural analysis of protein interfaces.

### Clustering of resolved interface structures

To quantify the diversity of experimentally resolved protein interfaces in the PDB, we developed the following workflow (Fig. 2a). We first extracted interacting chain pairs (dimers) from PDB biological assemblies using an *α*-carbon distance of *≤* 10Å. To reduce redundancy before all-by-all interface comparison, we clustered highly similar dimers by whole chain similarity. Strict thresholds (multimer TM-score *≥* 0.9, matched chain TM-scores *≥* 0.5) were used to avoid merging distinct interfaces. We then retained one representative structure from each dimer cluster, encoded their interfaces into 3Di, and clustered the representatives based on interface similarity. Finally, all members of each dimer cluster were assigned to the interface cluster of its representative (Fig. 2a). Starting from 3,121,961 protein dimers, whole-complex clustering reduced the dataset to 189,830 non-redundant dimer structures, whose interfaces were subsequently grouped into 77,167 interface clusters based on structural similarity (Table S7). The entire workflow required approximately six days: 21 hours on two L40S GPUs and 120 hours on a single 128-core node. To the best of our knowledge, this is the first comprehensive resource of experimentally determined protein interfaces clustered by interface structure.

**Fig. 2.**
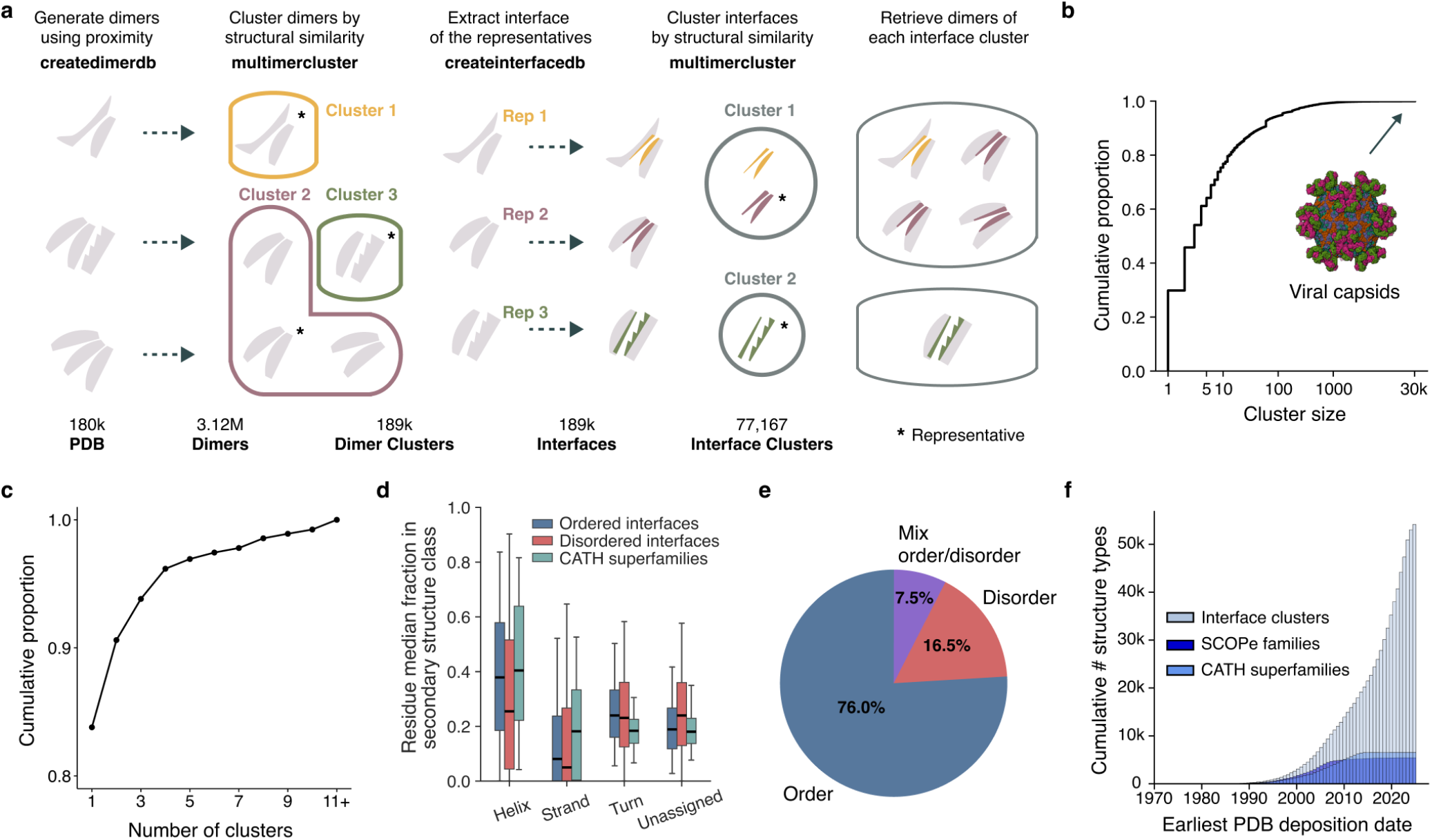
Clustering of resolved interface structures in the PDB. **(a)** Schematic showing the workflow for clustering all interface structures in the PDB. **(b)** Cumulative proportions of clusters from the PDB interface cluster resource by cluster size. The largest clusters consist mainly of viral capsid subunits (7YRH shown). **(c)** Cumulative proportions of NMR structures by the number of clusters obtained from clustering the corresponding NMR models. **(d)** Box whisker plots showing median fractions of ordered or disordered interface residues and CATH superfamily median fractions of residues per secondary structure type. Residue fractions are either computed over interface residues for each interface category or over entire folds for the fold category. Significances determined by mixed ANOVA, secondary structure × group interaction: *F* = 263.96, *P <* 0.001, Holm-corrected post hoc comparisons, all *P <* 0.001. **(e)** Pie chart displaying the proportions of interface clusters which contain only order-order interfaces, contain only interfaces with some degree of disorder (disorder-order or disorder-disorder), or contain some mix of order-order and disordered interfaces. **(f)** Growth of new CATH superfamilies or SCOPe families compared with newly observed interface clusters over time in the PDB.

The number of interface clusters and the proportion of singletons (one interface structure) strongly depend on the TM-score cutoff, with increasing cutoffs producing more singleton and fewer non-singleton clusters (Fig. S2a). At our chosen cutoff, 30% of interface clusters are singletons (Fig. 2b). Singleton and non-singleton clusters did not differ in disorder content, interface secondary structure, experimental method, or resolution (Fig. S2b-d). Compared with non-singleton interfaces, singleton interfaces were about 20 times more likely to originate from biological assemblies containing a single dimer (Fig. S2e). About 65% of clusters contained 2 to 100 members, while a few clusters contained thousands of interface structures, often from viral capsids (Fig. 2b). For 99.3% of non-singleton clusters, the average TM-score between the cluster representative and its members was at least 0.5 (Fig. S2f). The remaining low-scoring clusters often contained very small interfaces arising from incidental contacts between subunits of larger complexes and should be interpreted with caution (Fig. S2g).

Interfaces determined by different experimental methods rarely clustered together: only 8% of interface clusters contained structures resolved by more than one experimental method (Fig. S2h). This suggests that the range of experimental techniques available for structure resolution is useful for the discovery of new interface types. To assess to which extent conformational variability could lead to different interface cluster assignments, we clustered all NMR models from the same PDB entry. Only 16% of NMR entries yielded more than one interface cluster, suggesting that conformational flexibility at the interface, as captured by NMR, usually does not alter interface cluster assignment (Fig. 2c, S2i, Table S8). To characterize the structural properties of the interface clusters, we predicted disorder in the unbound state and classified interfaces as either ordered or disorder-mediated. Compared with protein folds, both ordered and disorder-mediated interfaces were enriched in turns, bends, and unassigned conformations, and were depleted in helices and strands (Fig. 2d, Table S9). More than half of all interface clusters contained only ordered interfaces (Fig. 2e). This contrasts with estimates that the majority of PPIs in eukaryotic interactomes are mediated by interfaces involving protein disorder (24), suggesting a substantial bias toward ordered interfaces in the PDB. New interface clusters continue to be observed even as superfamily discovery plateaus (Fig. 2f) (25), suggesting that the experimentally resolved interface space remains incomplete.

### Fold diversity in interface clusters

By focusing on interfaces rather than full chains, our approach can reveal shared interface geometries across proteins with different folds (26), providing potential examples of convergent interface evolution. To identify such cases, we annotated interfaces with CATH domains, using CATH’s hierarchical classification of protein structures (27). Among 13,725 clusters containing more than one CATH-annotated interface, only 721 (5%) contained more than one unique CATH domain pair (Fig. 3a). However, about 60% of these structurally diverse clusters contained folds from different CATH classes, indicating substantial divergence in overall structure despite similar interface geometry (Fig. 3b, Table S10). The number of unique gene ontology (GO) terms associated with proteins in a cluster correlated with the number of unique CATH domain pairs, suggesting that interface geometries carried by different folds are also used across diverse functions and cellular processes. Structurally diverse clusters also tended to have a higher fraction of helical residues at the interface (Fig. 3c). Although rare, these cases show that very different protein folds can arrive at similar local binding structures, providing candidate examples of convergent interface evolution.

**Fig. 3.**
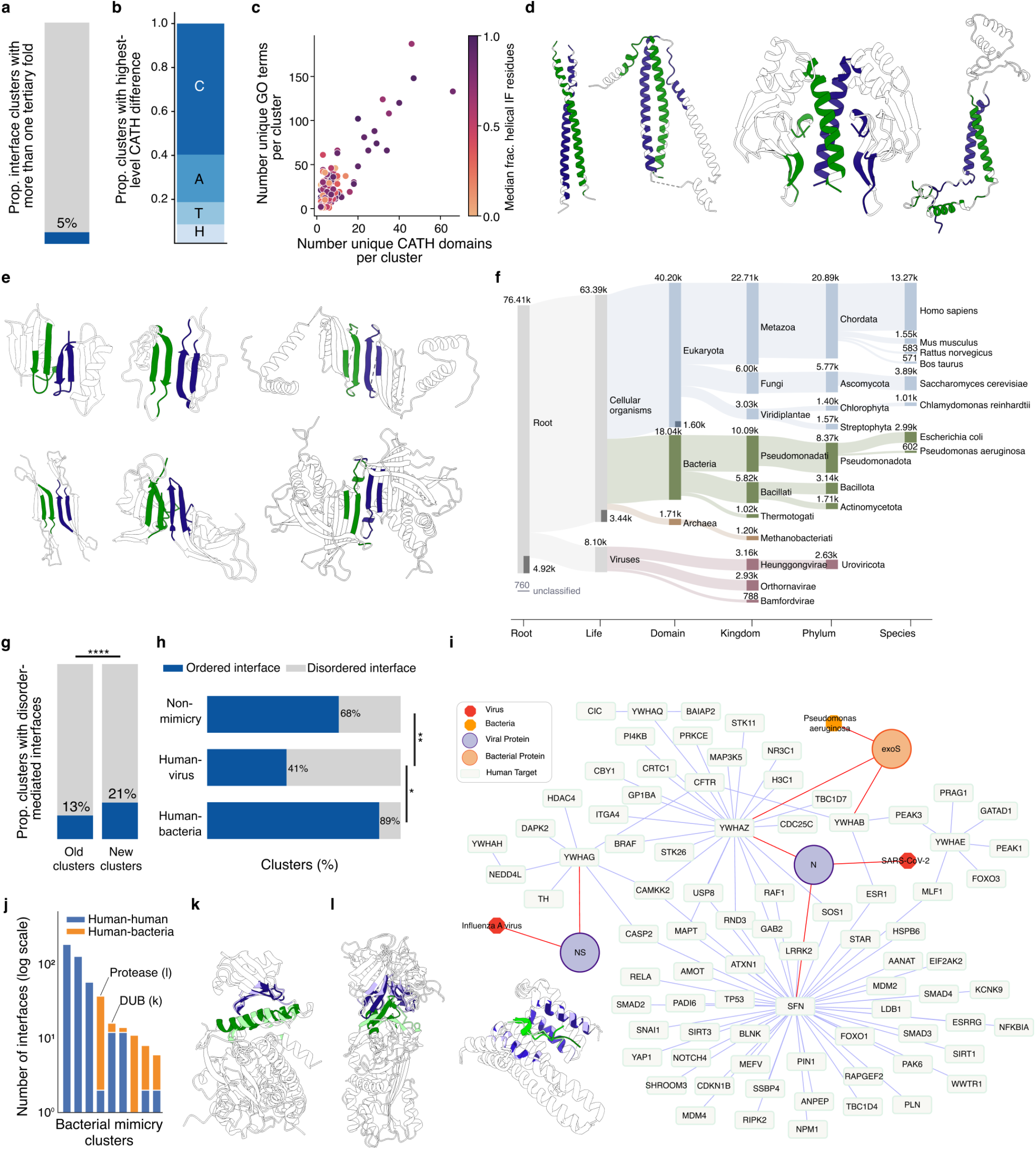
Structural diversity and evolution of interface clusters. **(a)** Proportion of clusters containing interfaces carried by more than one tertiary fold on at least one side of the interface. **(b)** Structurally diverse clusters grouped by the highest CATH level at which folds on either side of the interface diverge. **(c)** Correlation between structural and functional diversity across clusters. Point color indicates the fraction of helical interface residues. **(d)** Example interfaces from a structurally diverse cluster with a helical interface. Interface residues are highlighted in color. PDB IDs, from left to right: 5B83, 1J1D, 3DN7, 1V54. **(e)** Example interfaces from a structurally diverse cluster with a strand-like interface. Interface residues are highlighted in color. PDB IDs, from left to right: (top row) 2KWD, 4HSV, 1Q5V; (bottom row) 4FZQ, 3SB3, 4PN1. **(f)** Sankey plot of interface clusters grouped by their last common ancestor (LCA). **(g)** Proportion of evolutionarily old (LCA: cellular organisms) and human-only clusters containing disorder-mediated interfaces. Fisher’s exact test: statistic=0.59, p-value=5.7 *×* 10^−19^. **(h)** Proportion of predominantly disorder-mediated interfaces in HH-only clusters (non-mimicry) and clusters containing HH/HV or HH/HB interfaces. HH/HV cluster show higher proportion of disorder-mediated interfaces than non-mimicry clusters (OR = 0.33, p = 0.0008) and HH/HB cluster (OR = 11.5, p = 0.0226). **(i)** Network of structurally resolved PPIs from the only cluster containing HH, HV, and HB interfaces involving human 14-3-3 proteins (YWHAZ, YWHAG, SFN, YWHAB, YWHAE). Pathogen species of proteins are also indicated by edges. Inset: superimposed HH, HV, and HB interfaces (3UZD, 4O46, 2C23) aligned with Foldseek-Interface; viral and bacterial chains are shown in dark green and green, respectively. **(j)** Number of interface structures in each HH/HB cluster, split into HH and HB interfaces. Clusters illustrated in **k** and **l** are labelled. **(k)** Superimposed interfaces of human MINDY2 (light green, 7NPI) and an *O. tsutsugamushi* DUB (dark green, 6UPU), both bound to human ubiquitin (dark and light purple). **(l)** Superimposed interfaces of human PLAT (light purple) bound by SERPINE1 (light green, 5BRR) and human CTSG (dark purple) bound by *S. aureus* EapH2 (dark green, 6VTM).

One example of a structurally diverse cluster with a helical interface is represented by an Optineurin homodimer, which forms a coiled-coil interface (Fig. 3d). This cluster contains 443 members with 17 unique CATH domain pairs at the interface. Many of the CATH domain pairs involve the CATH topology 1.20.5: single alpha-helices involved in coiled-coils or other helix-helix interfaces; however, the cluster also contains interfaces from the mostly-beta CATH class (e.g., jelly rolls) and the few secondary structures CATH class (e.g., Cytochrome C oxidases) (Fig. 3d). GO enrichment analysis of this interface cluster revealed enrichment for proteins involved in aerobic respiration, specifically electron transport. A structurally diverse cluster with primarily strand interface structure is represented by a homodimer of the anti-tumor chemokine CXCL4L1, which dimerizes via beta sheets in the interface region (Fig. 3e). This cluster, with 727 members, contains 11 unique CATH domain pairs, about half from the mostly-beta CATH class and the other half from the mixed alpha-beta CATH class (Fig. 3e). This cluster shows enrichment for ribosomal subunits and proteins involved in lymphocyte activation, cell cycle and DNA metabolic process regulation, as well as transcription initiation. This analysis exemplifies the use of Foldseek-Interface and the interface cluster resource to identify and study cases of convergent interface evolution and the plasticity of different folds to evolve into common interface geometry.

### Evolution of interface structures

To categorize the PDB-derived interface clusters by evolutionary age, we retrieved the organisms of the interacting chains within each cluster and determined the last common ancestor (LCA) (Fig. 3f). We found that 4,917 clusters (6.4%) contain interfaces from both cellular organisms and viruses. A total of 3,440 clusters (4.4%) contain interfaces across all cellular organisms (Eukaryota, Bacteria, and Archaea) and were considered evolutionarily ‘old’. Conversely, 38,582 clusters (50%) were species-specific, about one third of which were human-only. We considered these clusters evolutionarily ‘young’, although some may be under-sampled in other phylogenetic groups. Comparing GO term frequencies between young and old clusters showed that young clusters are enriched in cellular signaling, immune cell differentiation, and organ development (Fig. S3a), functions associated with multicellularity and organismal complexity. Old clusters were instead enriched in iron ion transport, ATP synthesis, and biotin binding (Fig. S3a). In line with reports that disorder in proteomes increases with organism complexity (28), young clusters had a significantly higher proportion of disorder-mediated interfaces (21%) than old clusters (13%; Fig. 3g). This supports the idea that IDRs, including through short linear motifs, mediate transient regulatory and signaling events associated with multicellularity (24).

Pathogens can mimic host protein interaction interfaces to interfere with cellular machinery (29, 30). To identify potential cases of such mimicry, we searched for interface clusters containing both human-human (HH) and human-pathogen interfaces. After stringent filtering (see Methods), we identified 56 candidate mimicry clusters: nine containing HH and human-bacteria (HB) interfaces, 46 containing HH and human-virus (HV) interfaces, and one containing HH, HB, and HV interfaces (Table S11). HH/HB clusters largely contained ordered interfaces, while HH/HV clusters had a higher proportion of disorder-mediated interfaces than HH-only clusters (Fig. 3h). This is consistent with short linear motifs evolving more rapidly than protein folds and being more easily accommodated by small viral genomes, while bacterial proteomes contain relatively little disorder (28, 30). The mixed HH/HB/HV cluster consisted of phosphorylated motifs binding human 14-3-3 proteins (Fig. 3i) and captures the well-described hijacking of 14-3-3 proteins by pathogens to manipulate host trafficking and immune defense (29).

Focusing on bacterial mimicry, the nine HH/HB clusters varied substantially in size and fraction of HB interfaces (Fig. 3j), and five targeted the host ubiquitin-proteasome system. We identified four clusters with likely signs of convergent interface evolution, three of which, to the best of our knowledge, have not been described previously (31). One example showed highly similar interface geometry despite divergent folds between the human de-ubiquitinase (DUB) MINDY2 and the *Orientia tsutsugamushi* DUB (OTT_1962) (32, 33), both of which bind human ubiquitin (Fig. 3k). Similarly, the *Staphylococcus aureus* neutrophil serine protease inhibitor EapH2 and human SERPINE1 use similar interfaces to inhibit serine proteases despite their different folds (34, 35) (Fig. 3l), suggesting convergent evolution of bacterial proteins to target host proteases. The human targets of the 46 HH/HV clusters were enriched for apoptosis and chromatin-remodelling processes (Fig. S3b). The cluster spanning the largest number of viral species (11) corresponded to Bcl2-family-like viral folds targeting helical BH3 motifs in human proteins, thereby interfering with host apoptosis signaling (Fig. S3c,d). This represents a rare case of viral mimicry involving an entire fold (36). Another cluster spanning eight viral species captured the well-studied targeting of PDZ domain proteins involved in host cell polarity and cell-cell communication (Fig. S3e) (37). Together, these clusters enable systematic exploration of how bacterial and viral pathogens mimic host-like interface geometries sometimes using distinct protein folds.

### Novel interface discovery in predicted complexes

Tools such as AlphaFold are increasingly used to predict protein complex structures at scale. To assess whether AlphaFold can predict interface structures not represented during training, previous studies have largely used sequence- or fold-based splits or focused on ordered interfaces (11, 38). Using our PDB-derived interface clusters, we generated pre- and post-training benchmark datasets based directly on interface structural similarity. We randomly sampled 1,960 interface cluster representatives from clusters with interfaces deposited before or after the reported training cutoff of AlphaFold-Multimer (AF-MM) (39) and evaluated their prediction by AF-MM. Interfaces sampled from larger assemblies were predicted poorly overall (median DockQ 0.02; 83% of predictions below the 0.23 success threshold; Fig. S4c), likely because AF-MM cannot reconstruct the correct binding mode without the remaining subunits. We therefore restricted the comparison to 479 interfaces from assemblies containing a single interface, which resulted in overall higher prediction accuracies similar to previous reports (11, 38). AF-MM predicted post-training interfaces marginally worse on average than pre-training interfaces (Fig. S4a), with 75% accuracy among highly confident (ipTM *≥* 0.7) post-training predictions (Fig. 4a, Table S12). Accuracy was similar between order-order and disorder-order interfaces (Fig. S4b). However, disorder-disorder interfaces were more challenging to predict (Fig. S4b,c). Among the excluded larger complex interfaces the pre- versus post-training difference was significant (20.0% vs 15.4% with DockQ > 0.23, *P* = 0.023), but since both groups performed near the floor of the metric, we do not interpret this as evidence of reduced generalization.

**Fig. 4.**
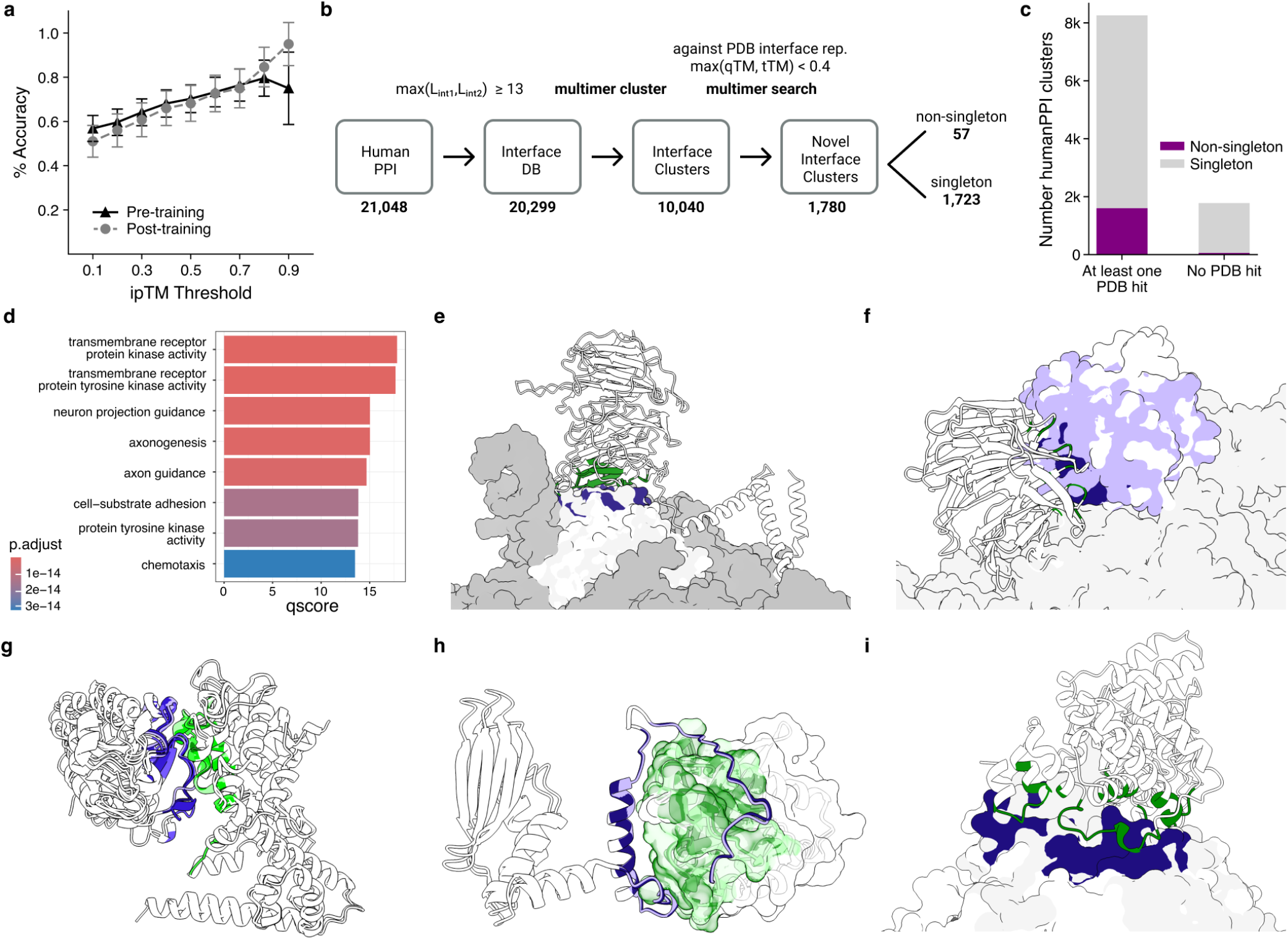
Evaluation of the prediction of novel interfaces by AlphaFold. **(a)** Accuracy of AlphaFold-Multimer predictions of interface clusters before and after the training cut-off date. Percent of top-ranked models with DockQ > 0.23 is calculated at increasing ipTM thresholds. **(b)** Schematic showing the search pipeline to identify putatively novel interface structures in the humanPPI dataset. **(c)** Stacked bar plot showing the number of clusters containing one or multiple interfaces and split by sharing similarity with at least one interface structure in the PDB or not. **(d)** Enriched GO terms in humanPPI interface clusters with putatively novel interface structures. **(e)** Superimposition of the predicted and putatively novel interface structure from humanPPI formed between RPS9A (purple) and EIF2A (green) and a structure from the ribosome including RPS9A (white, 7WTS). **(f)** Superimposition of a predicted interface from humanPPI between EIF6 (purple, surface) and PAK1IP1 (white, cartoon) and a resolved structure of the ribosome (6LQM) on EIF6 highlighting how the predicted binding of PAK1IP1 is compatible with EIF6 (white, surface) binding to other ribosomal subunits (grey, surface). **(g)** Superimposition of three predicted and putative novel interface structures from humanPPI between proteins from the ELMOD domain-containing protein family (green shades) and ARF-like GTPases (purple shades). **(h)** Superimposition of two predicted and putative novel interface structures from humanPPI between RABL2A/B (green shades) and CEP19 (purple shades). **(i)** Superimposition of a predicted and putatively novel interface structure from humanPPI formed between TBC1D23 (purple) and FHIP2A (green) and a resolved structure of the FHF complex involving FHIP1B (white, 8QAT).

Since these results suggest that AF-MM can predict interface structures not represented during training, we used Foldseek-Interface to search predicted protein complexes for putatively novel interface structures. We analyzed a published dataset of around 21,000 predicted binary human protein complexes generated using RosettaFold2 and AlphaFold2 (18), hereafter referred to as HumanPPI (Fig. 4b, S4d). We compared HumanPPI interfaces with each other and with PDB interface cluster representatives, resulting in 10,040 interface clusters. Of these, 1,780 contained only HumanPPI interfaces without structural similarity to any PDB interface cluster representative and were therefore considered putatively novel (Fig. 4b,c, Table S13). These clusters were generally smaller than clusters with similarity to resolved interfaces, with 97% being singletons (Fig. 4c, S4e). Disorder-mediated interfaces accounted for 24% of the putatively novel clusters, comparable to the full HumanPPI dataset but higher than in the PDB clusters (Fig. S4f). GO analysis showed an enrichment of transmembrane receptor kinases and neuron-related functions, including axonogenesis (Fig. 4d).

Manual inspection of 39 novel interface clusters (see Methods) revealed potential artifacts such as the docking of different partners or protein fragments to the same or multiple different surfaces (23 cases), poorly predicted residue-residue contacts (10 cases), artificial termini and exposed intramolecular contacts caused by fragmentation (2 cases), and contacts between extracellular and intracellular fragments (1 case). Some putatively novel interfaces also reflected conformational differences from resolved structures (3 cases, see Supplementary Information). Despite these caveats, six of the 39 inspected clusters contained putatively novel interfaces of particular interest. One predicted interface connects the alternative translation initiation factor EIF2A with the ribosomal subunit RPS9, positioning EIF2A on the ribosome surface (Fig. 4e). Another involves the eukaryotic translation initiation factor EIF6 and the PAK1 regulator PAK1IP1 (Fig. 4f). According to STRING, prior evidence for both interactions was weak. The well described interaction between TMIGD2 and HHLA2 involved in TCR-mediated T-cell activation (40), is predicted to occur between their extracellular Ig-like domains. All three interfaces belong to singleton clusters and are fully ordered. A cluster of four interfaces between ELMOD1/2/3 and ARF-like proteins (ARL6, ARL8A/B, ARFRP1) links proteins involved in GTPase regulation, membrane trafficking, and cilia function (41) (Fig. 4g). Two predicted interfaces between the centrosomal protein CEP19 and RABL2A/B also relate to cilia function (42) and involve an IDR of CEP19 predicted to wrap around RABL2A/B (Fig. 4h). Finally, two clustered interfaces between FHIP2A/B and TBC1D23 are consistent with evidence linking FHIP2A/B-containing FHF complexes to endosome-Golgi transport (43). The predicted interface is compatible with the recently resolved FHF complex containing FHIP1B, HOOK3, and AKTIP (44) (Fig. 4i). Together, these candidates span diverse cellular processes, folds, and IDRs, illustrating the potential of combining Foldseek-Interface with predicted complex structures to prioritize interfaces for further investigation.

### Web resource

Foldseek-Interface and the PDB interface cluster resource are accessible through two dedicated webservers. At https://search.foldseek.com/interface, users can query protein complexes containing at least two chains against prebuilt databases of PDB, AlphaFold Database homodimers (16), and HumanPPI interfaces. Structurally similar interfaces are returned with similarity scores, and an interactive viewer enables comparison at both the interface and full dimer level. The PDB interface cluster resource is available at https://interface.foldseek.com. Clusters can be searched by PDB identifier, UniProt accession, taxonomy, or a custom structure file. Each cluster provides summary information, taxonomic distribution and members at both the dimer and interface cluster level.

## Discussion

Already about two decades ago, scientists formulated the quest to systematically assess similarity between structurally resolved protein interaction interfaces regardless of overall fold similarity but tools were lacking to perform the required computations in reasonable time (12, 45). With the rapidly growing repertoire of protein complex structures in the PDB, this goal appeared even less in reach over time. Here, we developed an interface-specific structural encoding and clustering framework that makes systematic comparison of protein interfaces across the PDB computationally feasible. This enabled us to generate the first comprehensive protein interface cluster resource of resolved structures from the PDB. While developing the algorithms, we paid particular attention to the inclusion of small protein interaction interfaces, i.e. those mediated by short linear motifs, that are numerous in the PDB but often excluded in other large scale structure analysis efforts. With tailored benchmark datasets we carefully calibrated Foldseek-Interface to capture similarity between evolutionary related small interfaces while limiting the number of clusters consisting of accidental interface similarity. We found that the latter more often occurred for helical interfaces. Available annotations of our clusters, including various structure and confidence features, enable further post-hoc filtering depending on the desired analysis.

Both the cluster resource and Foldseek-Interface are made available as a web server, providing scientists with the ability to search for similar interface structure in the PDB and other resources of modeled complex structures. We expect Foldseek-Interface to become a new powerful tool for structural biologists, adding an intriguing new angle to protein complex structure analysis. Potential future applications include analyzing antibody binding and identifying off-targets to assess the specificity of designed protein binders. Similar to fold classification (27), Foldseek-Interface quantifies topological similarity between interfaces, disregarding biophysical properties that dictate binding affinity and specificity. Furthermore, interface residues are ordered from N- to C-terminus and are concatenated from both chains prior to 3Di conversion. In rare occasions, this may result in failure to detect interface similarity. Composite interfaces between two chains might also be classified as a new interface type despite consisting of sub-interface structures that share similarity with other clusters. Some of these limitations can be tackled in future developments of Foldseek-Interface. We used the interface cluster resource to generate AlphaFold pre- and post-training splits. Among interfaces from dimeric assemblies we observed no significant difference in prediction accuracy between pre- and post-training interface structures, in contrast to other reports (11, 38). We speculate that this is likely caused by a naive selection of interfaces from pre-training clusters disregarding filtering criteria employed by others such as interface size, presence of ligands, PTMs, and structure quality. Additionally, it is possible that splitting purely based on interface similarity is not stringent enough. Approaches that combine overall fold similarity and interface similarity might be more adequate for this task (11).

By clustering all protein interaction interfaces in the PDB based on similarity, we achieved a massive redundancy reduction from 3 million interfaces to 77,000 interface clusters, corresponding to a much larger number of structurally distinct interface clusters than previously estimated (12, 26). Akin to CATH for folds (27), we initially anticipated that interface clusters might organize into higher-order groups based purely on structural similarity. However, progressively lowering the similarity cutoff resulted in biologically meaningless groupings, suggesting that structural similarity alone does not reveal a clear hierarchy of interface types. Despite this, the resulting clusters proved useful for exploring remote interface relationships and evolution, identifying putatively novel interface types in large collections of predicted protein complexes, and revealing a strong bias toward ordered interfaces in the PDB. This bias suggests that the experimentally resolved interface universe remains substantially incomplete, particularly for disorder-mediated interactions, whose structural characterization will require approaches capable of capturing and comparing flexible and heterogeneous protein complexes.

## Methods

### createdimerdb module

We implemented a `createdimerdb` module to construct a dimer database from a Foldseek multimer database. For each input multimeric structure, all possible chain pairs are evaluated independently using a distance-based reciprocal contact criterion. A pair of chains is retained as one dimer entry if both chains contain at least four residues whose C-alpha atoms lie within 10 Å of a C-alpha atom in the other chain. This criterion is used only to determine whether the chain pair forms a valid dimeric pair; the output is the corresponding pair of chains. Therefore, a single multimeric structure can yield multiple dimer entries, and all qualifying chain pairs are collected into the final `dimerdb`.

### createinterfacedb module

We implemented a `createinterfacedb` module to construct an interface-level database from a Foldseek multimer database. If both chains contain at least four residues whose C-alpha atoms lie within 10 Å of a C-alpha atom in the other chain, these contact residues are defined as the interface and are written to the interface database. The amino acid sequence and C-alpha coordinate databases are subset to the extracted interface residues, whereas the 3Di states are recalculated for the interface structures to reflect their local structural environment. The resulting interface database enables downstream analyses focused specifically on protein–protein interaction regions rather than on full-chain dimeric structures.

### multimercluster module

We implemented a `multimercluster` module in Foldseek to cluster protein complexes based on structurally consistent multimer alignments. The module first uses `multimersearch` to align query and target complexes via multimer superposition, then groups them using `clust`. To account for the multifaceted nature of protein complex structure, we newly introduced four filtering criteria that collectively evaluate different aspects of structural similarity. First, we extended coverage to the multimer level. Instead of reporting coverage per indiviual chain, we aggregated over all aligned chains, weighting each by its length:

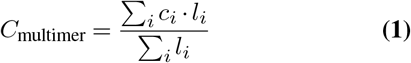

where *c*_*i*_ and *l*_*i*_ are the coverage and length of chain *i*, respectively. Coverage is computed separately for the query and target complexes. Second, the multimer TM-scores assess the global structural similarity of the full query and target complexes. Third, chain TM-scores are computed by applying the multimer-level superposition to each aligned chain pair individually, assessing per-chain structural similarity within the global alignment context. Fourth, an interface lDDT score is computed. Interface residues are defined respectively for the query and target structures as residues whose Cα atom lies within 10Å of the Cα atom of a residue in other chain. The lDDT score is then computed over these interface residues. Assignments satisfying all user-defined thresholds for these four criteria are then clustered by the clust module.

### Benchmark dataset

We created two benchmark datasets to assess the accuracy of Foldseek-Interface. Because no resource exists which clusters interface structures without using either of the methods we are benchmarking, we turned to sequence-based resources to generate datasets with clusters of interfaces which have similar sequence patterns, and therefore are presumed to have similar structures. The first dataset (Order-order) consists of interactions between ordered regions, or folded domains. This dataset was created by querying ProtCID (46) for all clusters related to a manually-curated dataset of 43 Pfam binding pairs which have a high-confidence domain-domain interface structure available in PDB (47). 30 of these Pfam binding pairs have associated ProtCID clusters; in total, the dataset contains 41 clusters and 654 interfaces. The ProtCID cluster assignment serves as the ground truth cluster assignment. The structure files for this dataset were fetched from a ProtCID data dump which was generated on February 1, 2024. The second dataset (Disorder-order) consists of interactions between an ordered region on one chain and a disordered region, mediated by a short linear binding motif (SLiM) on the other chain. This dataset was created by randomly selecting SLiM sequence classes from a data dump of the Motif Map of the Proteome (MoMaP) resource (https://slim-tools.org/momap/) which was generated on June 5, 2024. We select only MoMaP SLiM classes which are mapped to a motif class from the Eukaryotic Linear Motif (ELM) resource (48). The ELM class serves as the ground truth cluster assignment for this dataset. To select a balanced range of cluster sizes, we sampled an equal number of clusters from the second quartile of cluster sizes (2-6 members), the third quartile (6-15 members), and the fourth quartile (15+ members). In total, there are 58 clusters and 778 interfaces in the disorder-order dataset. Using the PDB ID and the two chain IDs associated with each interface, the structure files for the Disorder-order benchmark dataset were created by fetching the asymmetric unit PDB file from RCSB and using the Biopython v1.86 PDB module (49) to write a new structure file with only the two interacting chains. Interface size (number of interface residues per chain) was obtained from the Foldseek-Interface database files. Disorder content of the benchmark datasets was assessed by merging the disorder annotations for PDB interfaces to benchmark datasets based on PDB ID and chain IDs. See following sections for the disorder annotation of the PDB interfaces.

### Benchmarking Foldseek-Interface

We benchmarked Foldseek-Interface search by running pairwise all-by-all alignments with Foldseek-Interface commit e93bffb for each benchmark dataset separately. The alignment output contains two TM-scores, one normalized to the length of the query interface and one to the length of the target; these scores are averaged. Because the Foldseek-Multimer search parameters were optimized for full-length multimers, we needed to identify the optimal parameters to filter alignments between the shorter 3Di/AA sequences resulting from the Foldseek-Interface workflow. This filtering step reduces the number of interface pairs which get aligned and scored in the next step of the workflow, thereby reducing computational cost. We tested different pre-filtering methods (the default being k-mer match filtering) by adding flags –ungapped for an ungapped alignment pre-filter or `–exhaustive-search` to skip the pre-filtering step entirely and run all-by-all pairwise chain alignments. We also tested a range of E-values (0.001 to infinity) using the flag -e to see how accurately the E-values from the sequence alignments can assess interface chain similarity. These tests were performed on each benchmark dataset separately. We also tested these parameters by querying each benchmark dataset against a target dataset which included the benchmark dataset plus all PDB interfaces, to assess scalability and sensitivity in larger searches. Accuracy was determined by calculating the area under the ROC1 curve. ROC1 curves are generated by isolating all alignments for a given structure in the dataset, sorting these alignments in descending order of similarity, then calculating the fraction of members from that structure’s ground truth cluster (true positives; TP) which are sorted above the first member from another cluster (false positive; FP). These ROC1 curves are an apt assessment of alignment accuracy because they best reflect the most common real-world use-case of structure alignment methods. An area under the ROC1 curve of 1.0 would indicate a perfect agreement of alignment results with ground truth cluster assignments. When E-values proved insensitive for interfaces, necessitating overly-permissive thresholds, we turned to single-chain structure similarity as an alternative method of filtering single-chain alignments. We tested a range of TM-scores and lDDT values (0.1 to 0.9) using the flags `–tmscore-threshold` and `–lddt-threshold`. Tests for the TM-score and lDDT thresholds were performed on each benchmark dataset separately. Threshold efficacy was determined by calculating the sensitivity and specificity of each threshold compared to a baseline Foldseek-Interface search without any TM-score or lDDT threshold, with sensitivity calculated as *TP*_*thresh*_*/TP*_*baseline*_ and specificity calculated as (*FP*_*baseline*_ *™FP*_*thresh*_)*/FP*_*baseline*_. After parameter optimization, the final commands for benchmarking are:

~~~
foldseek createdimerdb
   --distance-threshold 10
   --min-interface-residues-perchain 4
foldseek createinterfacedb
   --distance-threshold 10
   --min-interface-residues-perchain 4
foldseek multimersearch
   --exhaustive-search -e 10000000
   --lddt-threshold 0.2
   --min-assigned-chains-ratio 1.0 --threads 1
~~~

We ran a similar pairwise all-by-all interface alignment with iAlign v1.1 (50) using the following commands, the first to test the iTM-score, the second to test the IS-score:

~~~
ialign.pl -s -n ave -e tm -o -minp 2 -mini 2 -dc 5
ialign.pl -s -n ave -o -minp 2 -mini 2 -dc 5
~~~

Accuracy for each benchmark method was assessed by drawing ROC1 curves and calculating the area under the ROC1 curve.

### Benchmarking Foldseek-Interface-Cluster

We benchmarked Foldseek-Interface-Cluster by clustering each benchmark dataset separately using foldseek commit e93bffb. First, we reduced redundancy in the full-length dimer structures with the following commands:

~~~
foldseek createdb \
foldseek createdimerdb
   --distance-threshold 10
   --min-interface-residues-perchain 4
foldseek multimercluster \
   --multimer-tm-threshold 0.9
   --chain-tm-threshold 0.5
   --cov-mode 0 --cluster-mode 0 -c 0
   --interface-lddt-threshold 0
~~~

We then performed interface clustering on the representatives of the dimer clustering with the following:

~~~
foldseek createinterfacedb
   --distance-threshold 10
   --min-interface-residues-perchain 4
foldseek multimercluster
   --multimer-tm-threshold 0.3
   --chain-tm-threshold 0
   --cov-mode 0 --cluster-mode 0 -c 0
   --interface-lddt-threshold 0
~~~

We also tested a multimer TM-score threshold of 0.4 and 0.5 for the interface clustering. Accuracy for Foldseek-Interface-Cluster was determined by calculating the Adjusted Rand Score, which is a measure of similarity in the way a set of items is clustered in a ground truth dataset versus the way the same set of items is clustered by a predictive method, which also accounts for the probability that the items could be randomly clustered correctly. The score ranges from -1 to 1, with 1 being a perfect replica of the ground truth clusters.

### Clustering PDB

Biological assembly files were downloaded from the PDBe (51) ftp server on March 13, 2025. Quaternary structures in the PDB are divided into dimers using `foldseek createdimerdb` with default parameters. The 3,121,961 dimer structures are clustered using foldseek commit e93bffb and the following command:

~~~
foldseek multimercluster
   --multimer-tm-threshold 0.9 -c 0
   --chain-tm-threshold 0.5 -cov-mode 0
   --interface-lddt-threshold 0 --cluster-mode 0
~~~

The interface region is retrieved from the representatives of 189,830 clusters using `foldseek createinterfacedb` with default parameters. Interface regions are then clustered with:

~~~
foldseek multimercluster
--multimer-tm-threshold 0.4 -c 0
--chain-tm-threshold 0 -cov-mode 0
 --interface-lddt-threshold 0 --cluster-mode 0
 --exhaustive-search 1 -e 10000000
 --lddt-threshold 0.2
~~~

To confirm the optimal multimer TM-score threshold during interface clustering, we re-ran the final interface clustering step with three different thresholds: 0.3, 0.4, and 0.5. We assessed the clustering accuracy by merging the ground truth cluster assignments of the benchmark structures based on PDB ID and chain IDs and computing the Adjusted Rand Score of the PDB interface cluster assignments to the ground truth cluster assignments for the benchmark structures.

### Cluster NMR structures

We identified 1,805 NMR structures that were included in our `dimerdb`. From these structures, we retrieved 5,135 dimers, of which 3,950 contained multiple NMR models. We then generated interface databases for these dimers and clustered the interface set of each NMR structure using the same parameters that were applied to the PDB and HumanPPI datasets. Overall, 83.79% of the NMR structures were grouped into a single cluster, whereas the most fragmented structure was divided into 25 clusters.

### Disorder analysis

Each interface chain was annotated with Uniprot IDs of the interacting chains using the EMBL-EBI SIFTS database (52). PDB residue seq IDs were mapped to Uniprot sequence positions using this database as well. Disorder at the interface was predicted using IUPred2A (53) (short option), by accessing the predicted residue-level disorder for a given UniProt ID. IUPred2A scores for each interface residue were extracted from the full-length canonical protein isoform predictions. An interface residue was considered disordered if the IUPred2A score was above 0.4. If at least 50% of the interface residues on a single chain were disordered, then that chain was considered to be disordered at the interface. Interfaces were then categorized as order-order, order-disorder, and disorder-disorder, based on the disorder status of the two chains in the interface. The order-disorder and disorder-disorder categories were then grouped into a single ‘disordered interface’ category.

### Secondary structure analysis

We compared the secondary structure composition of ordered and disordered interface clusters with the secondary structure composition of folds as defined by CATH homologous superfamily annotations in the PDB. Secondary structure assignments were extracted from pre-computed DSSP4 annotations in PDB-REDO (54) mmCIF files (downloaded from pdb-redo.eu on October 30, 2025) and classified into the categories helix (H, G, I, P), strand (E, B), turn/loop (T, S), and unassigned (55). The fractional abundance of each category was calculated for each CATH domain and aggregated at the homologous superfamily level using median values. Similarly, median fractional abundances were calculated for each ordered and disordered interface cluster. Secondary structure distributions across ordered interface clusters, disordered interface clusters, and CATH superfamilies were compared using a mixed ANOVA, followed by Holm-corrected two-sided pairwise t-tests.

### Growth of interface clusters over time

For each cluster, we created a list of PDB IDs contained in that cluster. We then annotated each PDB ID with its deposition date obtained from the ACCESSION DATE field of the wwPDB archive index file entries. We then took the earliest deposition date among all cluster members as the cluster’s date. To analyze the growth of protein folds over time, we downloaded the CATH v4.4 (56) domain list and made a list of PDB IDs for each CATH superfamily. We also downloaded the SCOPe release 2.08 (57), last updated January 6, 2023, and made a list of PDB IDs for each SCOPe family. Similar to the interface clusters, we then annotated PDB IDs with their release date and found the earliest release date for each CATH superfamily and for each SCOPe family.

### Singleton analysis

To check whether singleton clusters are driven by certain characteristics of the interface, we analyzed the prevalence of disorder and secondary structure elements using the annotations described above. We also fetched information about the experimental method and the resolution from the RCSB (58) GraphQL server based on PDB ID. This information was also used to analyze the experimental method composition of the interface clusters. To check whether the interface is from a dimer or from a larger complex, we counted the number of interfaces in our database for each biological assembly.

### Structure and function diversity analysis

We downloaded the CATH v4.4 (56) domain list and ‘seqreschopping’ domain boundaries dataset, which provides the PDB sequence residue ID start and end for each annotated CATH domain in the PDB. We merged the domains to the PDB interfaces based on PDB ID and chain ID, then retained only the domains whose boundaries overlap with at least one interface residue. To look for clusters with interface types carried by multiple tertiary folds, we first restricted that CATH annotations to those with exactly one domain on each interface chain, for ease of interpretation. We then excluded singleton clusters, clusters with majority protein-peptide or peptide-peptide interfaces (peptide is defined as < 20 resolved residues), clusters with an average representative-vs-member qTM-score < 0.5, and clusters which have 1 or fewer interfaces with a CATH annotation. After this filtering, 13,725 clusters remained. We label clusters as structurally diverse if they contain more than one CATH domain pair at the interface of different homologous superfamilies. We then further labeled clusters based on which CATH hierarchy level contains the divergence. We assessed functional diversity of the cluster by annotating each interface with a list of GO terms associated with the UniProt IDs involved in the interface, then counting the number of unique GO terms within a cluster. GO annotations were extracted from the UniProtKB Swiss-Prot (downloaded Jul 2025) and TrEMBL (downloaded Mar 2025) flat files by parsing each entry’s accession, GO term, and evidence code, and the two sets were concatenated into a single table. We performed functional enrichment on the structurally diverse cluster examples by submitting a list of the UniProt IDs associated with the most frequently occurring organism in the cluster to clusterProfiler (59).

### Taxonomic analysis

The taxonomy ID for each PDB chain was obtained from the mapping file of the Foldseek PDB database released in 2026, which postdates our PDB database(60). Cluster-level taxonomic assignments were performed using the MMseqs2 lca module (61), which reports the lowest common ancestor (LCA) for each cluster. For each interface cluster, the LCA was computed from the taxa of its member structures, including both the interface-cluster members and the members of their corresponding dimer clusters. Sankey diagrams were generated using the Metabuli-App(62).

For evolutionary analysis of interfaces, we labeled clusters whose LCA were ‘Cellular Organisms’ as ‘old’ interface clusters, and clusters whose LCA were Homo sapiens as ‘new’ interface clusters. Disorder analysis uses the IUPred2A annotations for the interfaces as described above. A cluster is labeled as ‘Disordered interfaces’ if the majority of the interfaces in the cluster are either order-disorder or disorder-disorder. We assessed functional enrichment in the old vs. new clusters by concatenating all GO annotations for all interfaces in the old clusters and for the new clusters respectively. For each GO term, we calculated the frequency with which it appears in the old clusters and the new clusters (the denominator is the total number of GO terms in the old and new clusters respectively). The fold enrichment is calculated by dividing the frequency of occurrence in the new clusters by the frequency of occurrence in the old clusters. We calculated significance by creating contingency tables containing the counts of that GO term and the counts of all other GO terms in the old clusters and the new clusters and performing a chi-squared test. All p-values are adjusted using Benjamin-Hochberg FDR correction.

### Pathogen mimicry analysis

We identified clusters of pathogen mimicry based on the PDB taxonomic annotation of chains at interfaces. Clusters containing both human-human and human-other (interspecies) interfaces were selected. These clusters were systematically filtered to exclude antibody interactions (cross-referenced annotation from SAbDab (63)), interfaces annotated as “IMMUNE SYSTEM” (according to PDB metadata), interactions involving *Escherichia coli*, and low-quality clusters with a repto-mem average cluster qTM score < 0.5. To eliminate clusters involving chimeric, synthetic, modified, or heavily mutated protein chains, we excluded any interface structure where one or more interacting chains lacked a valid UniProt annotation from the PDB. Clusters containing both human-human and human-pathogen (bacteria or virus) interfaces were defined as pathogen mimicry clusters, whereas clusters lacking human-pathogen interactions were classified as non-mimicry clusters.

All 14 human-bacteria clusters were manually checked to identify clusters that resulted from accidental interface similarity and were unlikely to represent cases of actual mimicry. Five such clusters were identified and removed. Manual checking of all 53 human-virus clusters was too laborious. Here, we first annotated all interacting chains with overlapping Pfam Hidden Markov Model matches. Interface residues were mapped to Pfam domains by intersecting their positions with Pfam domain boundaries. Pfam domain annotations of PDB structures were retrieved from RCSB on October 30, 2025. A Pfam domain was assigned to an interacting chain partner only if at least one residue participating in the structural interface fell within the defined sequence start-and-end boundaries of that domain. Clusters for which matched chains from the interfaces had divergent Pfam annotations were inspected manually resulting in the removal of 7 clusters. Of the remaining clusters, we compared the proportions of ordered and disordered interfaces (annotated as described in the *Disorder analysis*) among bacterial mimicry, viral mimicry, and non-mimicry clusters using pairwise two-sided Fisher’s exact tests with Benjamini-Hochberg FDR correction. Enrichment analysis of Gene Ontology biological processes for human targets from 46 viral mimicry clusters was performed using ShinyGO v0.85 (64). Network visualizations were generated using Cytoscape v3.10.4 (65).

### AlphaFold-Multimer benchmarking

We labeled PDB interface clusters as either ‘pre-training’ or ‘post-training’ based on the earliest PDB release date in the cluster (see above for annotation of release dates). If the earliest release date was after the training cut-off date for AF-MMv2.3 (September 30, 2021), we labeled the cluster as ‘post-training’. All other clusters were labeled ‘pre-training’. To sample interfaces from different categories, we labeled each cluster as ‘order-order,’ ‘order-disorder,’ or ‘disorder-disorder’ based on the majority interface type in the cluster (see ‘Disorder analysis’). We excluded clusters with majority peptide-peptide interfaces and with majority protein-protein interfaces with very small interface sizes (maximum interface chain length < 13 residues), then randomly sampled 300-400 clusters, depending on data availability, from each orderedness category from pre-training and post-training clusters.

For the cluster representatives, the PDB sequences (including unresolved residues) were retrieved from the RCSB (58) GraphQL server. Multiple sequence alignments were computed using MMseqs2 (commit 6f45232) (66) via `colabfold_search,` querying against the uniref30_2302 and colabfold_envdb_202108 databases at sensitivity 7, both downloaded from https://colabfold.mmseqs.com/. Structure prediction was then performed using ColabFold v1.5.5 (67) with 20 recycling iterations. We computed DockQ values for the predicted structures and analyzed the top-ranked models based on model confidence.

### Searching and clustering HumanPPI

We downloaded 21,048 predicted structures from the HumanPPI database (68) http://prodata.swmed.edu/humanPPI/, accessed on 29 September 2024. We retrieved 20,299 interfaces where at least one of the chains’ interface is longer than 12 residues. We clustered the interface database with the same parameters used to cluster the PDB. We also searched the interface database against the cluster representatives of the PDB interface clusters using `foldseek multimersearch` with default parameters. An interface was considered a hit if either its qTM or tTM score was above 0.4. We defined a HumanPPI cluster as a novel interface cluster if no cluster member had a hit against the PDB interface representatives.

### HumanPPI analysis

We annotated the HumanPPI interfaces with predicted disorder in a similar fashion as the PDB interfaces, fetching IUPred2A (53) (short) annotations for the UniProt IDs involved in the interactions, calculating the fraction of disordered interface residues (IUPred2A > 0.4), and labeling the interface chains as ordered or disordered. HumanPPI clusters are labeled as disordered if the majority of the interfaces in the cluster are either order-disorder or disorder-disorder. We performed functional enrichment of the novel interface clusters by running clusterProfiler (59) on the list of Uniprot IDs in the novel interface clusters. Foldseek-Interface search results return alignments and scores for interfaces which share similarity only on one chain, not both. We consider these partial matches to be no-hits, but we retained this information to better prioritize clusters for manual inspection. There were 38 clusters, entirely singletons, which did not have even a partial match to a PDB interface representative and which had an ipLDDT > 70; we manually inspected all of these clusters. We then filtered all other novel interface clusters for those which had an average ipLDDT > 70 and were non-singletons. We manually inspected most of these clusters. All structure images were rendered with ChimeraX v1.9 (69).

### Webserver

#### Search server

The Foldseek-Interface is integrated into the Foldseek search server as a new search mode. Users upload a protein complex in PDB or mmCIF format containing at least two chains and select one or more prebuilt interface databases. Three databases are provided: PDB, AlphaFold Database homodimers and HumanPPI. To reduce redundancy and search cost, PDB and AFDB homodimer databases were clustered at the dimer level using `foldseek multimercluster` with parameters `–multimer-tm-threshold 0.9 –chain-tm-threshold 0.5 –cov-mode 0 –cluster-mode 0 –interface-lddt-threshold 0`, reducing PDB from 3,121,961 to 189,830 representatives and AFDB homodimers from 1,757,658 to 634,776 representatives. HumanPPI was used as provided (20,953 structures). Only the representative of each dimer cluster was retained for searching. For every selected database, interfaces are searched with parameters `–multimer-tm-threshold 0.4 –cov-mode 0 -c 0 –exhaustive-search 1 -e 10000000 –lddt-threshold 0.20 –max-seqs 1000 –prefilter-mode 1 –cluster-search 0` and `–alignment-type` set to one of 3Di/AA, TM-align or LoLalign, as selected by the user. Result rows report TM-scores and sequence identities of interface regions. To enable comparison at both dimer and interface level, the corresponding structures are retrieved from a dimer database instead of interface database, and interface residues in the target are recollected by matching Cα coordinates returned by the alignment. The target chains are rebuilt from Cα to all-atom using PULCHRA (70), then the target structures are transformed using the translation and rotation matrix returned by Foldseek-Interface, and rendered with Mol*(71).

#### Cluster server

We developed a web server to explore the PDB interface clusters and the 3.1 million dimers they represent, building on the web framework developed for the AlphaFold Database cluster resource (72) and tailoring it for protein interface exploration. The server uses a REST-based client–server architecture with a Vue front-end and a Node.js back-end. Cluster and member metadata are stored in an SQLite database, and structures are streamed from Foldseek-derived databases through a C++ Node.js extension for fast read-in. We use WebAssembly-based versions of PULCHRA(70, 73) to restore full-atom structures from stored C-alpha traces and TM-align for pairwise alignment of interface regions between cluster members and their representatives, with structures rendered using Mol*(71). Taxonomic distributions of members are visualized as Sankey diagrams spanning superkingdom to genus. Structurally similar interfaces from HumanPPI are obtained by precomputed Foldseek-Interface searches of every cluster representative. Custom-structure queries are routed through the Foldseek-Interface search server, and the returned hits are mapped back to their cluster representatives.

## Data availability

Data can be searched at our dedicated web server https://interface.foldseek.com, is available for download at Zenodo https://doi.org/10.5281/zenodo.22040891 and as part of the supplementary material of this study.

## Code availability

Foldseek-Interface is free open-source software. The source code and ready-to-use binaries can be downloaded at github.com/steineggerlab/foldseek. The source code for the Foldseek-Interface search server is available at github.com/soedinglab/MMseqs2-App. The source code for the PDB interface cluster exploration server is available at github.com/rachelse/bfvd-web. Code written for data analysis is available at github.com/steineggerlab/foldseek-interface-analysis.

## Acknowledgements

We thank all members from the Steinegger and Luck lab as well as Pedro Beltrao for helpful discussions throughout the project. M.S. acknowledges support by the National Research Foundation of Korea (NRF) grants funded by the Korea government (MSIT) (RS-2024-00396026, RS2025-00438101, RS-2026-25549295) and the Novo Nordisk Foundation NNF24SA0092560. K.L. acknowledges funding from the Deutsche Forschungsgemeinschaft (DFG, German Research Foundation) project IDs 551068697 (financing J.M.S.), 449991970 and 464588647 - SFB1551, as well as funding from the European Union Horizon Europe research and innovation programme under the Marie Skłodowska-Curie grant agreement No 101227009 (ProtAIomics, financing H.S.). R.S.K. acknowledges support from the NRF (RS-2025-25405694).

## Author contributions

Authors are listed alphabetically. Conceptualization: K.L., M.S., J.M.S. Methodology: S.C., R.S.K, M.S., J.M.S. Software: S.C., R.S.K, M.S., J.M.S. Webserver: C.L.M.G., R.S.K. Formal analysis: S.C., H.S., J.M.S. Data Curation: K.L., J.M.S. Writing - Original Draft: S.C., R.S.K., K.L., H.S., M.S., J.M.S. Manuscript - Review and editing: S.C., C.L.M.G., R.S.K., K.L., M.S., J.M.S. Visualization: S.C., R.S.K., K.L., H.S., J.M.S., M.S. Supervision: K.L., M.S. Funding acquisition: K.L., M.S.

---

**Fig. S1.**
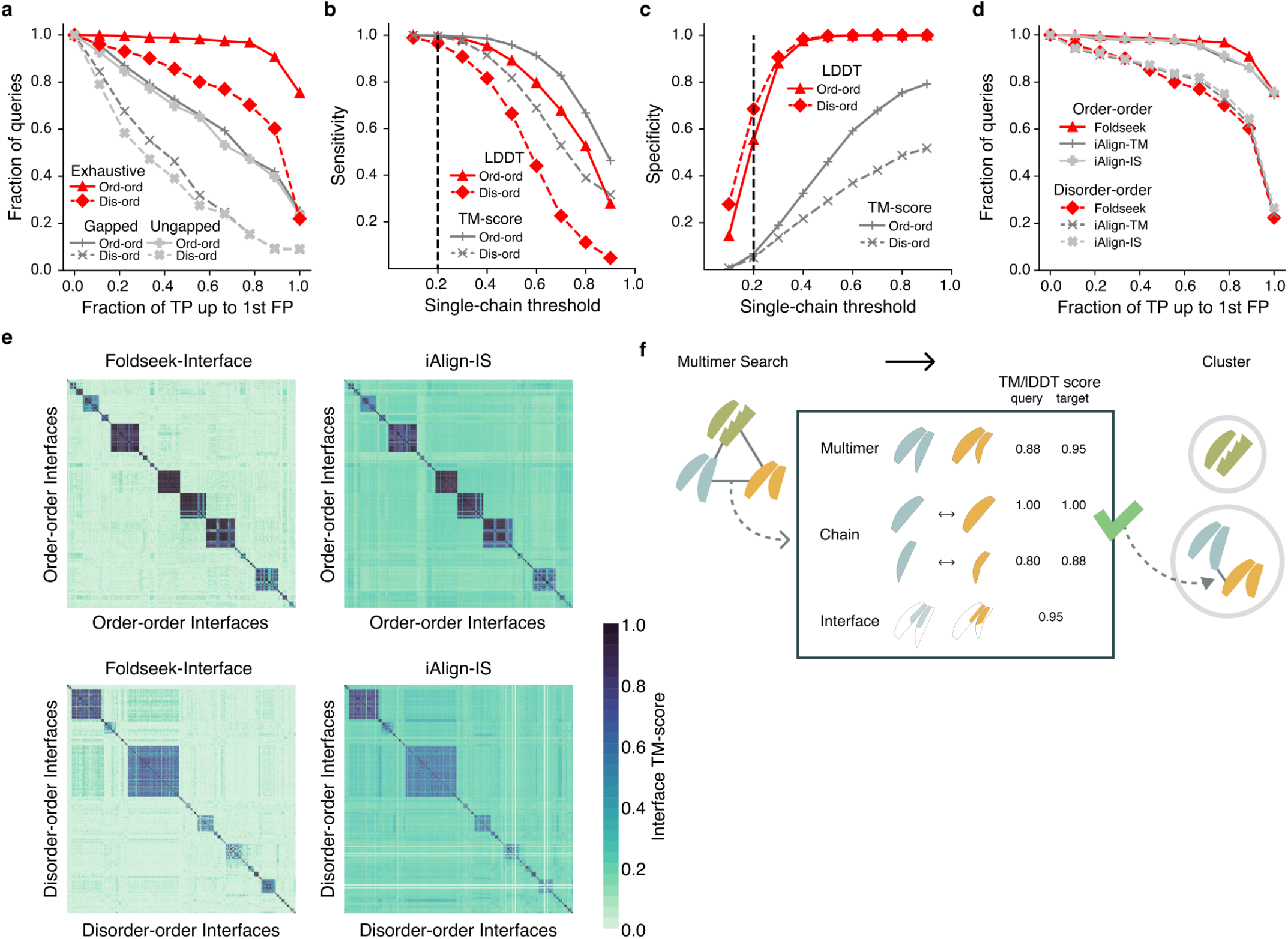
Benchmarking of Foldseek-Interface. **(a)** ROC1 curves of both benchmark datasets aligned to all 3.12M PDB interfaces plus benchmark structures to test sensitivity in large searches. Three filtering modes for the initial single-chain 3Di alignment step were tested with the exhaustive search performing best. **(b**,**c)** Sensitivity (b) and specificity (c) as a function of increasing single chain cutoffs for both benchmark datasets computed with respect to a baseline without any filters applied. Exhaustive search with an lDDT > 0.2 cutoff performed best. **(d)** ROC1 curves for Foldseek-Interface and iAlign (both scoring methods) for both benchmark datasets. **(e)** Heatmaps representing the all-by-all pairwise interface alignment scores for both benchmark datasets, for both Foldseek-Interface and iAlign (IS scoring method). **(f)** Schematic visualizing the three measures of similarity between two multimer structures, all three of which can be used as threshold to cluster interface or complex structures together.

**Fig. S2.**
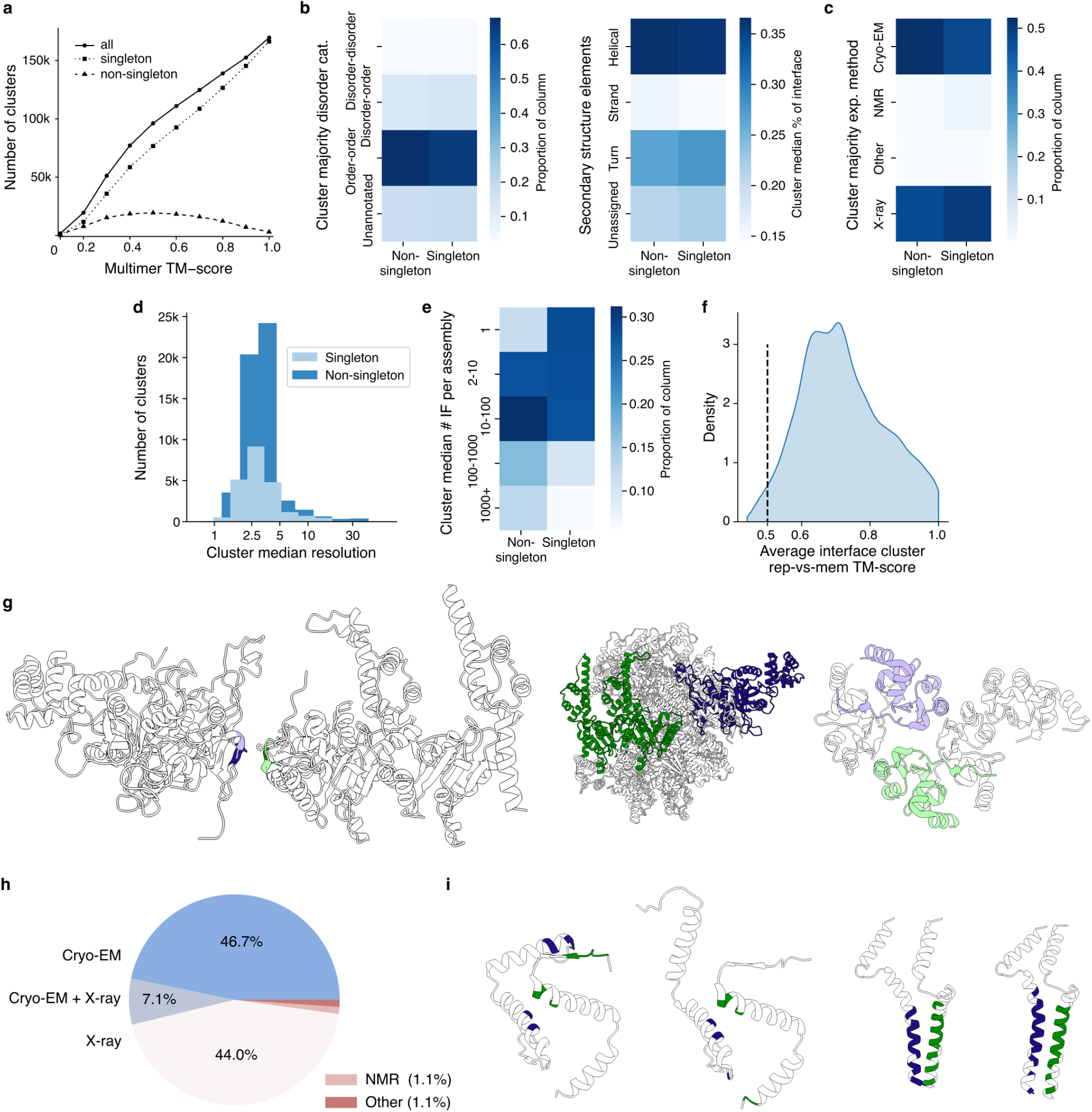
Characterization of PDB interface clusters. **(a)** Dotplot showing the number of clusters in total and split into singleton and non-singleton clusters as a function of increasing TM-score cutoffs. **(b)** Heatmaps comparing fractions of disorder and secondary structure in singleton vs. non-singletons clusters. **(c)** Heatmap comparing major experimental method of interfaces in singleton vs. non-singletons clusters. **(d)** Histogram showing the distribution of median experimental resolution of structures in singleton vs non-singleton clusters. **(e)** Heatmap comparing the fraction of interfaces in singleton vs non-singleton clusters split by the median number of interfaces in the PDB assemblies. **(f)** Density plot of the average interface TM-score of a cluster representative to each of its members across all the clusters. Dashed line represents the cut-off for determining if a cluster contains mostly low quality alignments. **(g)** Example of structures (8DAR A,G; 1OXQ A,B, assembly 6) from a low quality cluster that contains many small, secondary interfaces between two chains from larger complexes (left); illustration of the placement of the chains in the complexes (right). **(h)** Pie chart showing the fraction of clusters by experimental method used to resolve the interfaces in each cluster. **(i)** Examples of structures from clustering of NMR models. Both left and right panels are models which do not cluster together; left: 2NB1, chain B, C; right: 2HYN, chain A, E.

**Fig. S3.**
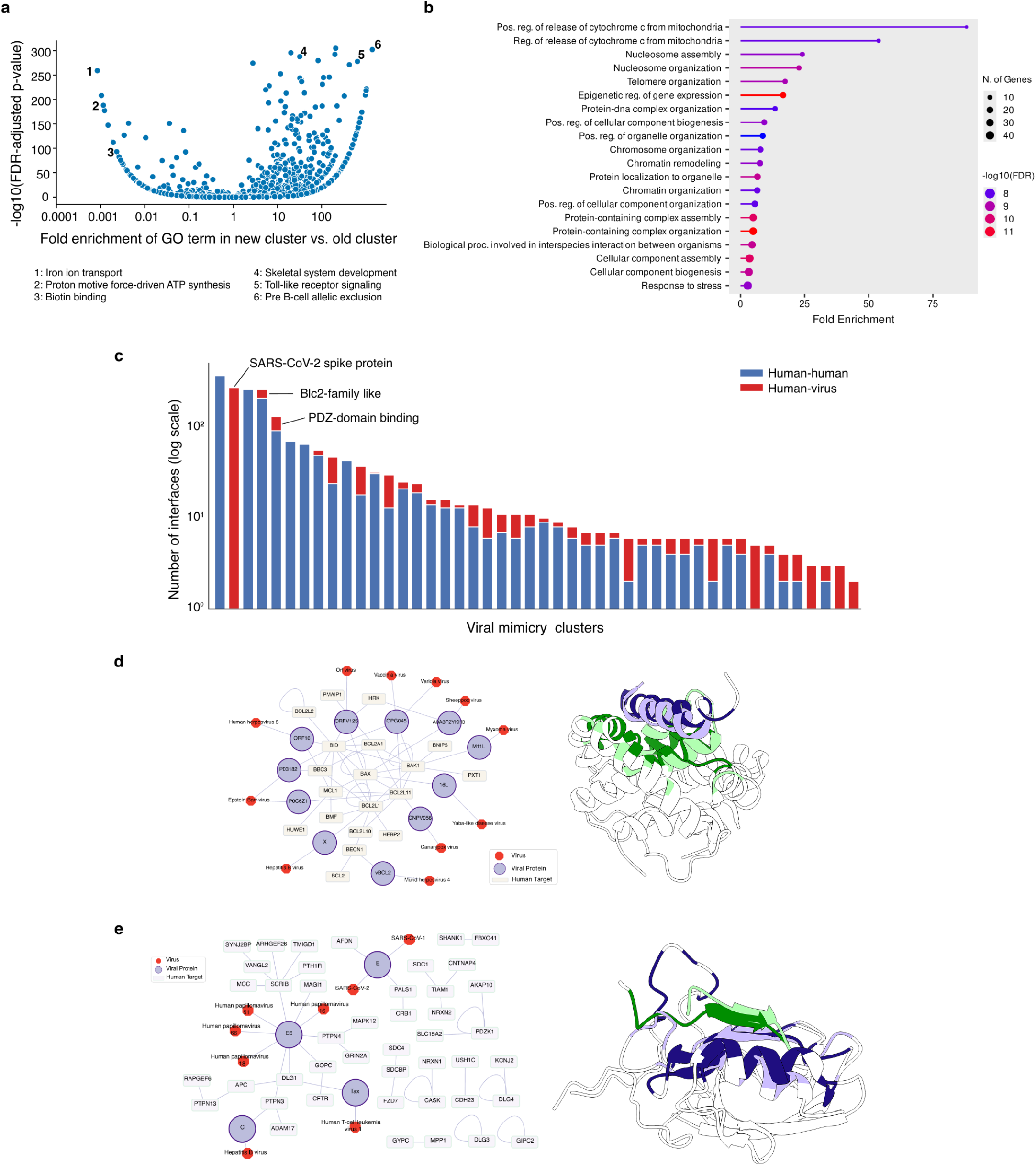
Aspects of interface evolution. **(a)** Volcano plot showing enrichment of GO terms in new clusters vs. old clusters, with fold-enrichment on x-axis and Benjamini Hochberg-adjusted p-values on the y-axis. Examples of highly-enriched terms in old and new clusters are labeled. **(b)** Top enriched GO terms for human proteins targeted by viral proteins in the identified HH/HV clusters. **(c)** HH/HV clusters ordered by their number of interfaces from largest to smallest indicating for each cluster the fraction of interfaces that are HH or HV. Three HH/HV clusters of interest are labelled, two of which are also shown in **d** and **e. (d)** Network of HH and HV interfaces in the HH/HV cluster of the Bcl2 family-like structures. Interfaces between proteins are shown as edges, proteins are shown as rectangle or round nodes. Species of viral proteins are shown as heptagonal shapes (left). Superimposition of a representative HV (dark purple/green, 2JBY, green: viral protein) and HH (light purple/green, 3I1H) interface from this cluster (right). **(e)** Network of HH and HV interfaces in the HH/HV cluster of the PDZ domain family, style as in **d** (left). Superimposition of a representative HV (dark purple/green, 2KPL, green: viral protein) and HH (light purple/green, 3RL7) interface from this cluster (right).

**Fig. S4.**
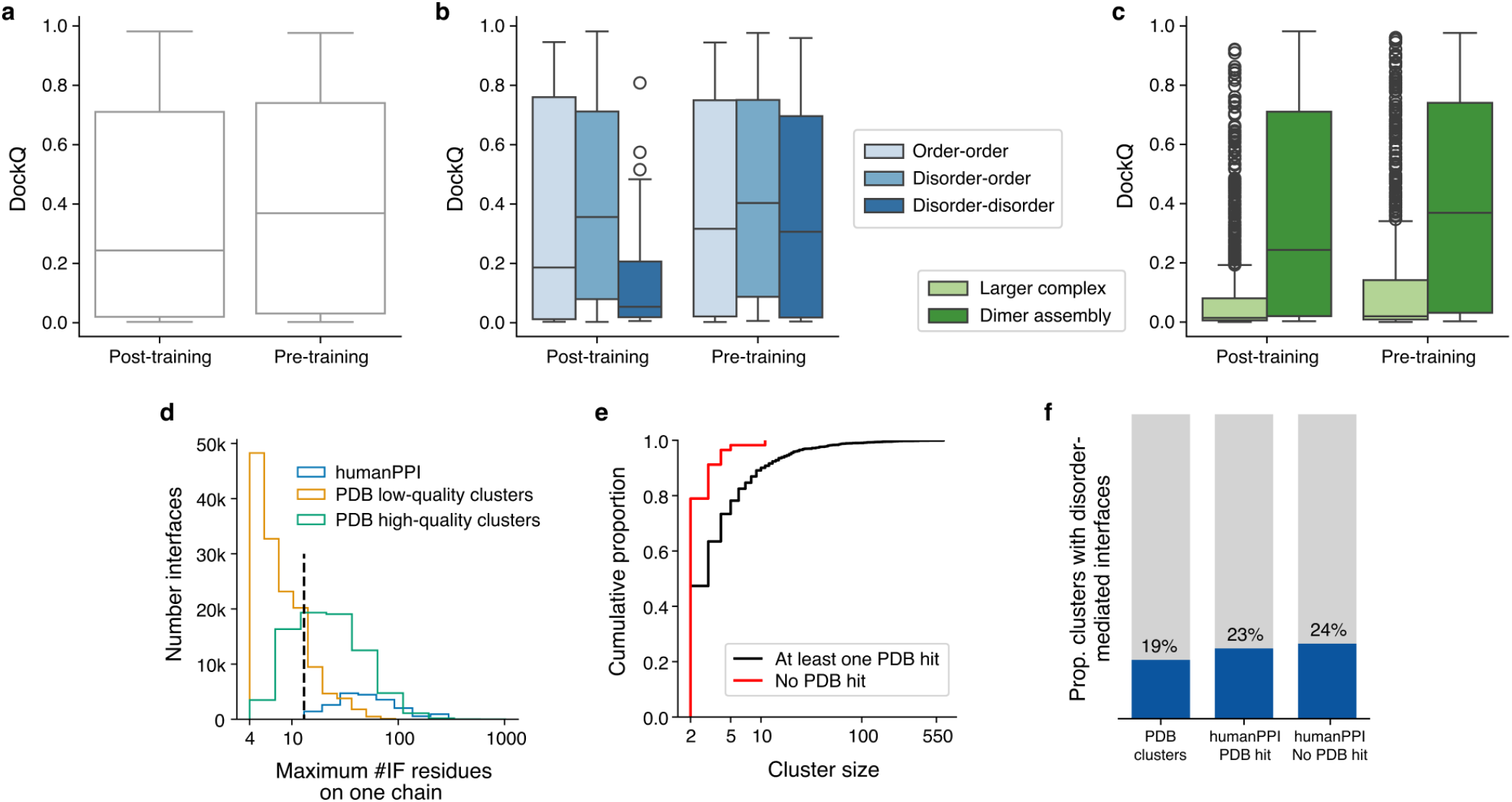
Prediction of novel interface types by AlphaFold-Multimer. **(a)** Box whisker plots showing the distribution of DockQ scores for top-ranked predictions of interface representatives from clusters with earliest deposition date before the AF-MM training cut-off date (pre-training) and after the cut-off date (post-training). Difference is not significant based on Student’s T-test (statistic=1.26, p-value=0.21, df=477). **(b)** Distribution of DockQ scores split by disorder content of the interface and pre- vs post-training (see a). Sample sizes, from left to right: n=91, n=81, n=22, n=95, n=95, n=95. **(c)** Distribution of DockQ scores split by whether the interface is from an assembly with only one interface (dimer assembly) or from an assembly with more than one interface (larger complex) and pre- vs post-training (see a). Sample sizes, from left to right: n=810, n=194, n=671, n=285. **(d)** Distribution of interface size for humanPPI, interfaces from low quality PDB clusters (average cluster interface TM-score *<* 0.5), and interfaces from other PDB clusters. Interface size is determined by the maximum number of interface residues on a single chain in the interface. Dashed line represents an interface size of 12, which is used as a cut-off to exclude humanPPI interfaces from analysis. **(e)** Cumulative proportion of humanPPI clusters by size and split by sharing similarity with at least one PDB cluster representative or not. **(f)** Proportion of clusters with disorder-mediated interfaces for PDB clusters, humanPPI clusters with similarity to PDB cluster representatives and humanPPI clusters without similarity to resolved interface representatives in the PDB.

## References

1. Orengo, C. A. & Thornton, J. M. Protein families and their evolution-a structural perspective. Annu. Rev. Biochem. 74, 867–900 (2005).

2. Mizianty, M. J., Peng, Z. & Kurgan, L. MFDp2: Accurate predictor of disorder in proteins by fusion of disorder probabilities, content and profiles: Accurate predictor of disorder in proteins by fusion of disorder probabilities, content and profiles. Intrinsically Disord. Proteins 1, e24428 (2013).

3. Van Roey, K. et al. Short linear motifs: ubiquitous and functionally diverse protein interaction modules directing cell regulation. Chem. Rev. 114, 6733–6778 (2014).

4. Jumper, J. et al. Highly accurate protein structure prediction with AlphaFold. Nature 596, 583–589 (2021).

5. van Kempen, M. et al. Fast and accurate protein structure search with foldseek. Nat. Biotechnol. 42, 243–246 (2024).

6. Barrio-Hernandez, I. et al. Clustering predicted structures at the scale of the known protein universe. Nature 622, 637–645 (2023).

7. Henschel, A., Kim, W. K. & Schroeder, M. Equivalent binding sites reveal convergently evolved interaction motifs. Bioinformatics 22, 550–555 (2006).

8. Cukuroglu, E., Gursoy, A., Nussinov, R. & Keskin, O. Non-redundant unique interface structures as templates for modeling protein interactions. PLoS One 9, e86738 (2014).

9. Winter, C., Henschel, A., Tuukkanen, A. & Schroeder, M. Protein interactions in 3D: from interface evolution to drug discovery. J. Struct. Biol. 179, 347–358 (2012).

10. Vallat, B. et al. RCSB protein data bank: Delivering integrative structures alongside experimental structures and computed structure models. Nucleic Acids Res. 54, D489–D498 (2026).

11. Kovtun, D. et al. PINDER: The protein interaction dataset and evaluation resource. bioRxiv 2024.07.17.603980 (2024).

12. Kim, W. K., Henschel, A., Winter, C. & Schroeder, M. The many faces of protein-protein interactions: A compendium of interface geometry. PLoS Comput. Biol. 2, e124 (2006).

13. Dapkūnas, J., Timinskas, A., Olechnovič, K., Tomkuvienė, M. & Venclovas, PPI3D: a web server for searching, analyzing and modeling protein-protein, protein-peptide and protein-nucleic acid interactions. Nucleic Acids Res. 52, W264–W271 (2024).

14. Kim, H., Kim, R. S., Mirdita, M., Yoon, J. & Steinegger, M. Structural motif search across the protein universe with folddisco. Nat. Biotechnol. 1–5 (2026).

15. Gao, M. & Skolnick, J. iAlign: a method for the structural comparison of protein-protein interfaces. Bioinformatics 26, 2259–2265 (2010).

16. Han, Y. et al. AlphaFold database expands to proteome-scale quaternary structures. bioRxiv 2026.03.27.714458 (2026).

17. Qi, X. et al. Atlas of predicted protein complex structures across kingdoms. Nat. Commun. 17, 4397 (2026).

18. Zhang, J. et al. Predicting protein-protein interactions in the human proteome. Science 390, eadt1630 (2025).

19. Humphreys, I. R. et al. Computed structures of core eukaryotic protein complexes. Science 374, eabm4805 (2021).

20. Kim, W. et al. Rapid and sensitive protein complex alignment with foldseek-multimer. Nat. Methods 22, 469–472 (2025).

21. Zhang, Y. & Skolnick, J. Scoring function for automated assessment of protein structure template quality. Proteins 57, 702–710 (2004).

22. Xu, Q. & Dunbrack, R. L., Jr. ProtCID: a data resource for structural information on protein interactions. Nat. Commun. 11, 711 (2020).

23. Kumar, M. et al. ELM-the eukaryotic linear motif resource-2024 update. Nucleic Acids Res. 52, D442–D455 (2024).

24. Tompa, P., Davey, N. E., Gibson, T. J. & Babu, M. M. A million peptide motifs for the molecular biologist. Mol. Cell 55, 161–169 (2014).

25. Levitt, M. Nature of the protein universe. Proc. Natl. Acad. Sci. U. S. A. 106, 11079–11084 (2009).

26. Gao, M. & Skolnick, J. Structural space of protein-protein interfaces is degenerate, close to complete, and highly connected. Proc. Natl. Acad. Sci. U. S. A. 107, 22517–22522 (2010).

27. Waman, V. P. et al. CATH v4.4: major expansion of CATH by experimental and predicted structural data. Nucleic Acids Res. 53, D348–D355 (2025).

28. Xue, B., Dunker, A. K. & Uversky, V. N. Orderly order in protein intrinsic disorder distribution: disorder in 3500 proteomes from viruses and the three domains of life. J. Biomol. Struct. Dyn. 30, 137–149 (2012).

29. Via, A., Uyar, B., Brun, C. & Zanzoni, A. How pathogens use linear motifs to perturb host cell networks. Trends Biochem. Sci. 40, 36–48 (2015).

30. Davey, N. E., Travé, G. & Gibson, T. J. How viruses hijack cell regulation. Trends Biochem. Sci. 36, 159–169 (2011).

31. Couves, E. C. et al. Structural basis for membrane attack complex inhibition by CD59. Nat. Commun. 14, 890 (2023).

32. Abdul Rehman, S. A. et al. Mechanism of activation and regulation of deubiquitinase activity in MINDY1 and MINDY2. Mol. Cell 81, 4176–4190.e6 (2021).

33. Berk, J. M. et al. A deubiquitylase with an unusually high-affinity ubiquitin-binding domain from the scrub typhus pathogen orientia tsutsugamushi. Nat. Commun. 11, 2343 (2020).

34. Gong, L. et al. Crystal structure of the michaelis complex between tissue-type plasminogen activator and plasminogen activators inhibitor-1. J. Biol. Chem. 290, 25795–25804 (2015).

35. Herdendorf, T. J., Stapels, D. A. C., Rooijakkers, S. H. M. & Geisbrecht, B. V. Local structural plasticity of the staphylococcus aureus evasion protein EapH1 enables engagement with multiple neutrophil serine proteases. J. Biol. Chem. 295, 7753–7762 (2020).

36. Kvansakul, M. & Hinds, M. G. Structural biology of the bcl-2 family and its mimicry by viral proteins. Cell Death Dis. 4, e909 (2013).

37. Castaño-Rodriguez, C., Honrubia, J. M., Gutiérrez-Álvarez, J., Sola, I. & Enjuanes, L. Viral PDZ binding motifs influence cell behavior through the interaction with cellular proteins containing PDZ domains. Methods Mol. Biol. 2256, 217–236 (2021).

38. Guan, L. & Keating, A. E. Training bias and sequence alignments shape protein-peptide docking by AlphaFold and related methods. Protein Sci. 34, e70331 (2025).

39. Evans, R. et al. Protein complex prediction with AlphaFold-multimer. bioRxiv 2021.10.04.463034 (2021).

40. Zhu, Y. et al. B7-H5 costimulates human T cells via CD28H. Nat. Commun. 4, 2043 (2013).

41. Turn, R. E. et al. The ARF GAPs ELMOD1 and ELMOD3 act at the golgi and cilia to regulate ciliogenesis and ciliary protein traffic. Mol. Biol. Cell 33, ar13 (2022).

42. Nishijima, Y. et al. RABL2 interacts with the intraflagellar transport-B complex and CEP19 and participates in ciliary assembly. Mol. Biol. Cell 28, 1652–1666 (2017).

43. Christensen, J. R. et al. Cytoplasmic dynein-1 cargo diversity is mediated by the combinatorial assembly of FTS-hook-FHIP complexes. Elife 10, e74538 (2021).

44. Abid Ali, F. et al. KIF1C activates and extends dynein movement through the FHF cargo adapter. Nat. Struct. Mol. Biol. 32, 756–766 (2025).

45. Keskin, O., Tsai, C.-J., Wolfson, H. & Nussinov, R. A new, structurally nonredundant, diverse data set of protein-protein interfaces and its implications. Protein Sci. 13, 1043–1055 (2004).

## References

46. Xu, Q. & Dunbrack, R. L., Jr. ProtCID: a data resource for structural information on protein interactions. Nat. Commun. 11, 711 (2020).

47. Geist, J. L., Lee, C. Y., Strom, J. M., de Jesús Naveja, J. & Luck, K. Generation of a high confidence set of domain-domain interface types to guide protein complex structure predictions by AlphaFold. Bioinformatics 40, btae482 (2024).

48. Kumar, M. et al. ELM-the eukaryotic linear motif resource-2024 update. Nucleic Acids Res. 52, D442–D455 (2024).

49. Cock, P. J. A. et al. Biopython: freely available python tools for computational molecular biology and bioinformatics. Bioinformatics 25, 1422–1423 (2009).

50. Gao, M. & Skolnick, J. iAlign: a method for the structural comparison of protein-protein interfaces Bioinformatics 26, 2259–2265 (2010).

51. Armstrong, D. R. et al. PDBe: improved findability of macromolecular structure data in the PDB. Nucleic Acids Res. 48, D335–D343 (2020).

52. Dana, J. M. et al. SIFTS: updated structure integration with function, taxonomy and sequences resource allows 40-fold increase in coverage of structure-based annotations for proteins. Nucleic Acids Res. 47, D482–D489 (2019).

53. Mészáros, B., Erdos, G. & Dosztányi, Z. IUPred2A: context-dependent prediction of protein disorder as a function of redox state and protein binding. Nucleic Acids Res. 46, W329–W337 (2018).

54. Joosten, R. P., Long, F., Murshudov, G. N. & Perrakis, A. The PDB_REDO server for macromolecular structure model optimization. IUCrJ 1, 213–220 (2014).

55. Hekkelman, M. L., Salmoral, D., Perrakis, A. & Joosten, R. P. DSSP 4: FAIR annotation of protein secondary structure. Protein Sci. 34, e70208 (2025).

56. Sillitoe, I. et al. CATH: increased structural coverage of functional space. Nucleic Acids Res. 49, D266–D273 (2021).

57. Fox, N. K., Brenner, S. E. & Chandonia, J.-M. SCOPe: Structural classification of proteins–extended, integrating SCOP and ASTRAL data and classification of new structures. Nucleic Acids Res. 42, D304–9 (2014).

58. Vallat, B. et al. RCSB protein data bank: Delivering integrative structures alongside experimental structures and computed structure models. Nucleic Acids Res. 54, D489–D498 (2026).

59. Yu, G., Wang, L.-G., Han, Y. & He, Q.-Y. clusterProfiler: an R package for comparing biological themes among gene clusters. OMICS 16, 284–287 (2012).

60. Han, Y. et al. AlphaFold database expands to proteome-scale quaternary structures. bioRxiv 2026.03.27.714458 (2026).

61. Mirdita, M., Steinegger, M., Breitwieser, F., Söding, J. & Levy Karin, E. Fast and sensitive taxonomic assignment to metagenomic contigs. Bioinformatics (2021).

62. Lee, S. J., Kim, J., Mirdita, M. & Steinegger, M. Easy and interactive taxonomic profiling with metabuli app. bioRxiv, 2025. 03. 10. 642298 (2025).

63. Dunbar, J. et al. SAbDab: the structural antibody database. Nucleic Acids Res. 42, D1140–6 (2014).

64. Ge, S. X., Jung, D. & Yao, R. ShinyGO: a graphical gene-set enrichment tool for animals and plants. Bioinformatics 36, 2628–2629 (2020).

65. Shannon, P. et al. Cytoscape: a software environment for integrated models of biomolecular interaction networks. Genome Res. 13, 2498–2504 (2003).

66. Steinegger, M. & Söding, J. MMseqs2 enables sensitive protein sequence searching for the analysis of massive data sets. Nat. Biotechnol. 35, 1026–1028 (2017).

67. Mirdita, M. et al. ColabFold: making protein folding accessible to all. Nat. Methods 19, 679–682 (2022).

68. Humphreys, I. R. et al. Computed structures of core eukaryotic protein complexes. Science 374, eabm4805 (2021).

69. Meng, E. C. et al. UCSF ChimeraX: Tools for structure building and analysis. Protein Sci. 32, e4792 (2023).

70. Rotkiewicz, P. & Skolnick, J. Fast procedure for reconstruction of full-atom protein models from reduced representations. J. Comput. Chem. 29, 1460–1465 (2008).

71. Sehnal, D. et al. Mol* viewer: modern web app for 3D visualization and analysis of large biomolecular structures. Nucleic Acids Res. 49, W431–W437 (2021).

72. Barrio-Hernandez, I. et al. Clustering predicted structures at the scale of the known protein universe. Nature 622, 637–645 (2023).

73. van Kempen, M. et al. Fast and accurate protein structure search with foldseek. Nat. Biotechnol. 42, 243–246 (2024).

