## Supplementary information for "Foldseek-Interface reveals a protein interface universe far from complete"

### Notes on manually checked putatively novel predicted interfaces in humanPPI

1. P49639 - Q9UQ80: Homeobox protein Hox-A1 (HOXA1) and Proliferation-associated protein 2G4 (PA2G4) humanPPI entry link
    a. Disorder-disorder, 52 interface residues
    b. Highest-scoring match bears little resemblance to the predicted interface (TM-scores < 0.2)
    c. Interaction NOT found in PPI databases, no PDB homologues
    d. *Common cellular compartment:* Nucleus
    e. *Common biological process keywords*: Transcription, Transcription regulation
    f. *Why is this one interesting?* Predicted interface is a disordered tail of HOXA1 docked into a large pocket on the PA2G4 domain. Classified as Disorder-disorder, which could possibly be due to the presence of some disordered loops in the pocket, or could be an IUPred2A inaccuracy. It appears to be Disorder-order at least. Could not find evidence of known PA2G4 binding motifs in MoMaP.
    g. The motif in HOXA1 is a known LIG_HOMEOBOX motif (elm.eu.org) and according to structure 1B72 is binding to homeobox domains; it could also be that the pocket on the PA2G4 where the HOXA1 motif is docked into binds RNA, it is a quite wide pocket and the domain is known to bind nucleic acids; the motif does not make good contacts with the pocket residues
2. A0A1W2PQJ7 - Q5H9L4: RNA polymerase II subunit A C-terminal domain phosphatase SSU72 like protein 1 (SSU72L1) and Transcription initiation factor TFIID subunit 7-like (TAF7L) humanPPI entry link
    a. Disorder-order, 22 interface residues
    b. No interface matches with PDB
    c. Interaction NOT found in PPI databases, no PDB homologues
    d. *Common cellular compartment*: Nucleus
    e. *Common biological process keywords:* None (Individual keywords include mRNA processing, Protein phosphatase, Developmental protein, Differentiation, Transcription)
    f. *Why is this one interesting?* This is the cluster representative of the only non-singleton in the potentially novel interfaces. The other member is A0A1W2PQJ5 - Q5H9L4 (a paralogue of SSU72L1: SSU72L3). The predicted interface is rather small and may be biophysically unconvincing (needs further inspection), but the predicted interface is similar with the two paralogues. I'm unsure how biologically interesting this is, given that RNA Pol II has several structures in PDB. But perhaps not this subunit or not interfacing with TIF.
    g. SSU72L1 is a phosphatase known to dephosphorylate the CTD of RNAP2, TAF7L might be a spermatogenesis-specific factor of TFIID; TAF7L is predicted to bind with polyPro loop to the domain of SSU72L1, this polyPro loop is also confidently predicted in the same conformation in monomer model of TAF7L, so will certainly bind something, the loop is docked against two exposed Phe residues on SSU72L1; models with other partner proteins of TAF7L that are not

paralogs of SSU72L1 are not very convincing and do not use this polyPro loop; SSU72L1 belongs to the SSU72 family of phosphatases -> there is a structure 4IMI that shows how a member of this family binds to a peptide from the CTD -> the active site is elsewhere from where the polyPro peptide of TAF7L is predicted to bind; rerunning the protein pair on AlphaFoldServer produces the same interface but of moderate confidence; I found a homologous structure 4RGW between TAF7 (paralog of TAF7L) and TAF1 -> superimposition of 4RGW with this structural model shows that the polyPro loop is involved in binding to TAF1, the fold of TAF1 and TAF7L is highly similar but how TAF1 and SSU72L1 bind to the polyPro loop is very different; I conclude that this structural model is likely wrong

3. P02458 - Q6ZMI3: Collagen alpha-1(II) chain (COL2A1) and Gliomedin (GLDN) [humanPPI entry link]
    a. Disorder-order, 47 interface residues
    b. Highest-scoring match in PDB shows no similarity
    c. Interaction found in STRING, no PDB homologues
    d. *Common cellular compartment:* Extracellular matrix/Secreted
    e. *Common biological process keywords:* None (one protein has no keywords), individual keywords are Developmental protein, Differentiation, Neurogenesis
    f. *Why is this one interesting?* The interface has a unique topology. Part of the large IDR of COL2A1 is bundled up in beta-hairpins and binds to the GLDN Olfactomedin-like domain. Unsure how convincing this interface is, but it could potentially be novel as I see no structure in PDB of Olfactomedin-like domain bound to anything. Unsure if the COL2A1 IDR is known to bind in this fashion to anything else. PDB structures for COL2A1 seem to focus only on its single folded domain.
    g. The residue-residue contacts between both protein chains do not look very convincing. The region of COL2A1 that is predicted to bind to the beta-propeller repeat domain of GLDN is in an interesting beta-sheet structure that is not of high confidence (light blue for pLDDT) in the monomer model of COL2A1 but it is in the same conformation. For both proteins fragments are used in this model. I looked at other complex models involving GLDN and found that other top scoring models all have some beta-only fold that is docked onto this surface of the propeller domain but all these folds are very different. COL2A1 is known to function in cartiloginous tissue while GLDN has a well studied role in brain development. I am unfortunately rather skeptical about this interface.
4. Q13084 - Q8TAE8: Large ribosomal subunit protein bL28m (MRPL28) and Large ribosomal subunit protein mL64 (GADD45GIP1) [humanPPI entry link]
    a. Disorder-disorder, 15 interface residues
    b. Highest-scoring match: 2cmn A-4,A-5 (qTM: 0.395, tTM: 0.25)
    c. Interaction is included in BIOGRID and STRING
    d. Has many PDB homologues
    e. This interface is quite small and based on the protein names, I am assuming these are two subunits of a large complex (ribosome). Given that we know there

are multiple ribosomal structures in the PDB, I would guess that this particular interface is not very interesting to study further. The reason it does not match any PDB interface representative could be due to small conformational differences in the disordered loop, or it could be an incorrectly predicted binding site.

5. P12270 - Q8NHY2: Nucleoprotein TPR (TPR) and E3 ubiquitin-protein ligase COP1 (COP1) humanPPI entry link
    a. Disorder-order, 48 interface residues
    b. Highest-scoring match: 7vgg A,B (qTM: 0.34, tTM: 0.24)
    c. 7vgg interface and predicted model interface both contain an instance of a motif which matches COP1 WD40 degron regex (from MoMaP) bound to COP1 WD40 domain. The similarity is not reflected in the TM-scores because the motif backbone is in a slightly different conformation
    d. The predicted instance of this motif is not annotated in MoMaP
    e. Interaction is found in STRING
6. P60866 - Q86VM9: Small ribosomal subunit protein uS10 (RPS20) and Zinc finger CCCH domain-containing protein 18 (ZC3H18) humanPPI entry link
    a. Disorder-order, 24 interface residues
    b. Highest-scoring match: 8i0t A,K (qTM: 0.36, tTM: 0.09)
    c. Interaction is found in BIOGRID
    d. Similar to the other ribosomal interface above, I'm not sure if this is interesting since it involves a ribosomal subunit. The interface does seem to involve the zinc-finger domain of ZC3H18 so that could be of interest. The highest-scoring match is a Zinc finger domain bound to a spliceosome subunit.
    e. not very convinced by the interface, the small Zn-finger domain binds a loop of RPS20 but it looks strange, all other models with ZC3H18 and other partner proteins do not involve the Zn finger domain
7. O43747 - P63010: AP-1 complex subunit gamma-1 (AP1G1) and AP-2 complex subunit beta (AP2B1) humanPPI entry link *Note: the interface in question is O43747_S2__P63010_S1, which is not the interface which autopopulates in the webserver
    a. Disorder-order, 18 interface residues
    b. Highest-scoring match bears little resemblance
    c. Interaction is found in BIOGRID and STRING, several PDB homologues
    d. I doubt the novelty of this interface because it is a helical motif of AP1G1 docked into the ARM-like domain of AP2B1. This seems valid (the monomer AFDB structure for AP1G1 even has this helical motif in the IDR), but given that there are several known ARM-binding motifs, I wonder if this is a novel structure. I have not thoroughly checked if the motif sequence matches any known motif regex patterns, and I also have not checked if the predicted pocket on ARM-like domain is the same as previously resolved structures. So there could still be novelty here and is worth reviewing.
    e. Adapter protein complexes (1-4) exist of 4 subunits, they are responsible for sorting cargo into clarthin-coated vesicles
       AP1: consists of two large adaptins (gamma and beta), one medium adaptin (mu)

and one small adaptin (sigma): Gamma -> AP1G1, beta: AP1B1, mu -> AP1M1/2, sigma -> AP1S1/2/3
AP2: consists of AP2A1/2, AP2B1, AP2M1, AP2S1
Structure 6CM9 shows how the AP-1 assembles -> AP1G1 and AP1B1 bind to each other with their Arm-repeat domains like predicted by HumanPPI but it is structure O43747_S1_P630101_S1, within the curvature that is formed by each Arm-repeat domain, the AP2M1/2 subunits bind -> this is where this short motif is predicted to bind of AP1G1 onto the ARM-repeat domain of AP2B1 -> I would think that this interface is usually not accessible because covered by AP-1 mu subunits, because of all of this observations I would think that this prediction is unlikely to be true

8. Q02543 - Q96EY4: Large ribosomal subunit protein eL20 (RPL18A) and Translation machinery-associated protein 16 (TMA16) humanPPI entry link
    a. Disorder-order, 18 interface residues
    b. Highest-scoring match: 5o31 m,n (qTM: 0.22, tTM: 0.06)
    c. Interaction is found in STRING, some PDB homologues
    d. Similar to the other ribosomal interfaces above, I wonder how interesting this would be to study.
    e. according to structure 6LSS TMA16 is part of the pre-60S ribosomal subunit and the interaction interface of TMA16 that is predicted here to bind to RPL18A is in the structure bound by RPL21; there are also other structural models with TMA16 in humanPPI and they are predicted to bind different folds at the same interface of TMA16, so overall not very convincing
9. Q12788 - Q86WX3: Transducin beta-like protein 3 (TBL3) and Active regulator of SIRT1 (RPS19BP1) humanPPI entry link
    a. Disorder-order, 74 interface residues
    b. Highest-scoring match: 2p91 C,F (qTM: 0.24, tTM: 0.20)
    c. Predicted model motif structure does not match the motif structure of the highest-scoring match, although the domain is the same.
    d. Helical motif “docked” on the rim of WD40 repeat domain of TBL3. There is a motif listed in MoMaP which binds to TBL WD40 domain but the predicted motif here does not match the regex. Nor does it match the regex for any of the WD40 binding motifs I found in MoMaP (at least, not by quick glance). The structure of the predicted motif also does not match the resolved structure for the TBL WD40 binding motif.
    e. This interaction is included in STRING, so perhaps the predicted binding region is incorrect?
10. P30405 - P55854: Peptidyl-prolyl cit-trans isomerase F, mitochondrial (PPIF) and Small ubiquitin-related modifier 3 (SUMO3) humanPPI entry link
    a. Disorder-order, 45 interface residues
    b. Highest-scoring match: 2x2a A,B (qTM: 0.22, tTM: 0.12)
    c. Interaction NOT found in PPI databases, no PDB homologues
    d. *Common cellular compartment*: None
    e. *Common biological process keywords*: None

    f. the binding face of PPIF appears to be the same face it uses to dimerize in PDB 2x2a.

11. O75717 - Q92771: WD repeat and HMG-box DNA-binding protein 1 (WDHD1) and Putative ATP-dependent RNA helicase DDX12 (DDX12P) [humanPPI entry link] *Note: the interface in question is O75717_S1__Q92771_S2, which is not the structure which automatically populates in the web server
    a. Disorder-order, 24 interface residues
    b. Highest-scoring match: 3nzp A,B (qTM: 0.25, tTM: 0.02)
    c. Interaction NOT found in PPI databases, no PDB homologues
    d. *Common cellular compartment:* Nucleus
    e. *Common biological process keywords:* DNA-binding
    f. I think this may be incorrect because the interface consists of the C-terminal tail of WDHD1 inserted into the C-terminal helicase domain of DDX12P. However, this 'tail' of WDHD1 is an artifact of the segmentation. The full protein is ~3x as long as the modeled segment. In reality, I'm not sure this interface would be physically available considering the orientation of the IDR in the binding pocket.

http://prodata.swmed.edu/humanPPI/results/O94985_Q63HQ2 (S2_S2) 121 IFres
For EGFLAM no structure resolved with binding partners, just monomers
No structure exists for CLSTN1
EGFLAM functions at retinal photoreceptor synapse, mediates transsynaptic interactions, extracellular
CLSTN1 is a postsynaptic adhesion molecule mediating synapse formation, intracellular at membranes
PPI with high confidence in STRING, PPI got two different high quality predictions with different fragments, which is a problem, other than that hard to tell if these are true or wrong predictions, they involve laminin, fibronectin, EGF-like domains

http://prodata.swmed.edu/humanPPI/results/Q15678_Q92626 (S2_S1) 115 IFres biology of both pair does not match well (PTPN14,PXDN)
PXDN rather extracellular
There are 4 different models with fragments of these proteins that are high confident -> this starts making me skeptical about the accuracy of these predictions

http://prodata.swmed.edu/humanPPI/results/Q2VWP7_Q8NFZ4 (S2_S0) 85 IFres looks interesting but maybe not novel -> need to compare to 5XEQ; the interface in the predicted model is mediated by approximately the same residues as in 5xeq, and the secondary structure elements are the same. The conformations/orientation are different, hence the missed match, conclusion: not novel

http://prodata.swmed.edu/humanPPI/results/P08648_P78324 (S2_S0) 81 IFres
Interaction between ITGA5 (integrin a5) and SIRPA (Tyr-protein phosphatase non-receptor type substrate 1)
There are two different confident predictions with different fragments -> one involves the Ig like

folds in both domains, and one involved the WD repeat domain in ITGA5 and a disordered region/motif in SIRPA;
These predictions with the Ig-like domains are hard to interpret, the same is true for predictions with WHD repeat domains and both are different but predicted for the same protein pair

http://prodata.swmed.edu/humanPPI/results/Q6EMK4_Q9Y219 (S0_S1) 72 IFres
Interaction between VASN (vasolin) and JAG2 (jagged-2), both function in growth signaling (TGF and Notch, respectively)
JAG2 has a lot of EGF-like folds, VASN a large LRR-repeat domain and there are two models that predict in different ways how these domains might bind each other
PPI is also in STRING
Other than that, not sure about the prediction

http://prodata.swmed.edu/humanPPI/results/P40818_P60484 (S2_S0) 70 IFres no structure displayed in humanPPI, had to download both models
Interaction between USP8 and PTEN -> both models show very different interfaces, hard to tell, in the PDB no useful structures resolved with other binding partners for both proteins to further evaluate

http://prodata.swmed.edu/humanPPI/results/P62502_Q6UWW0 (S0_S0) 68 IFres
This is a kind of homoderimerization between two lipocalin domains from two different lipocalin proteins, LCN6 and LCN15. There is a study (PMID: **26346541**) that describes different ways by which lipocalins can dimerize. One of the structures they show has some similarity to ours. So, in principle, this structural model could be true but functionally is not very exciting.

http://prodata.swmed.edu/humanPPI/results/P46781_Q9BY44 (S1_S0) 67 IFres
Interaction between the ribosomal subunit RPS9 and the alternative translation initiation factor EIF2A (this factor is not part of the canonical EIF2 complex, which also has a subunit called EIF2A but the gene/protein name for this one is EIF2S1). There is some weak evidence of functional association between EIF2A and RPS9. It is not known how EIF2A interacts with the ribosome to initiate translation but it is different from the EIF2 complex which needs GTP while EIF2A binds in a codon-dependent way. What is super interesting is that most of the surface of RPS9 interacts with other ribosome subunits including rRNA but the short bit that is accessible is predicted to interact with EIF2A. Superimposition of the PDB 7WTS with this structural model shows this nicely. The interface itself does not show a lot of specific contacts but it is quite a large interface. EIF2A is not predicted to bind any other protein in the humanPPI dataset. RPS9 has a lot of other structural models with other proteins. There is a review about knowledge of alternative translation initiation factors: doi:10.1002/wrna.1833

http://prodata.swmed.edu/humanPPI/results/O75473_P17181 (S1_S0) 62 IFres
Interaction between LGR5 and IFNAR1. LGR5 is an atypical G-protein coupled receptor that functions in Wnt signaling while INFAR1 is an interferon alpha/beta receptor. The structural model predicts an interaction between the extracellular Leucine-rich repeat domain of LGR5 and the extracellular fibronectin domains of IFNAR1. IFNAR1 is known to function in JAK/STAT

signaling. The LRR domain of LGR5 is known to bind r-spondins, there is also a structure resolved (4BST), which shows how the LRR domain and r-spondin form a heteromer. The contact sites overlap a lot with the predicted interfaces of the LRR domain with the fibronectin domains. Given the distinct biology of both proteins and this interface being known to bind something else, I don’t give much credibility to this prediction.

http://prodata.swmed.edu/humanPPI/results/P19022_P48960 (S1_S1) 49 IFres
Interaction between CDH2 and ADGRE5. CDH2 is Cadherin-2 functioning in mediating cell-cell contacts in neuronal context. ADGRE5 not well studied, supposed to function in cell adhesion processes after leukocyte activation (adhesion G-protein coupled receptor E5). Predicted interface is between an EGF-like domain in the extracellular region of CDH2 and an LRRNT domain in the extracellular region of ADGRE5. I am not sure about all the predictions of interactions between extracellular domains… The predicted interface is also quite small despite being between two folded domains.

http://prodata.swmed.edu/humanPPI/results/O75054_Q9HBX8 (S2_S1) 48 IFres
Interaction between IGSF3 (immunoglobulin superfamily member 3) and LGR6 (leucine-rich repeat containing G protein coupled receptor 6). Again a predicted interface between two extracellular regions, Ig-like domains in IGSF3 and the LRR domain in LGR6. Ig domain might bind to LRR domain where it usually binds R-spondins (see earlier notes). I think this is a very questionable interface prediction.

http://prodata.swmed.edu/humanPPI/results/Q13241_Q15485 (S0_S0) 44 IFres
Interaction between KLRD1 (natural killer cells antigen CD94) and FCN2 (Ficolin-2). KLRD1 functions in immune signaling in self-non-self recognition. FCN2 acts as a pattern recognition receptor that initiates the lectin pathway of the complement system, functioning in phagocytosis and breakdown of pathogens. Predicted interface is very very small, between two domains of each protein. Quite questionable prediction.

http://prodata.swmed.edu/humanPPI/results/P23434_P48728 (S0_S0) 44 IFres
Interaction between GCSH (Glycine cleavage system H protein) and AMT (aminomethyltransferase, T protein). Both proteins are part of the mitochondrial glycine cleavage system complex composed of GCSH, AMT, GLDC, DLD. There is a structure resolved with GCSH bound to LIAS (mitochondrial lipoyl synthase) (8UGO) which makes an important modification to GCSH for it to function properly. The predicted interface between GCSH and AMT overlaps much with the interface between GCSH and AMT. However, I would not for this reason necessarily conclude that this predicted interface is wrong. To further investigate this I modelled the whole GCS complex to see how AF would arrange all four proteins and if the same interface between GCSH and AMT would be repredicted. No structure of the complex seems resolved so far. It turns out that in the AF model of the whole complex GLDC is docked onto the same surface of GCSH like AMT and LIAS. AMT is not predicted to be in contact with GCSH anymore. Thus, the prediction is rather questionable. AF also did not predict well the GCS complex. Maybe important cofactors are missing.

http://prodata.swmed.edu/humanPPI/results/Q8IUA0_Q9H114 (S0_S0) 42 IFres
Interaction between WFDC8 and CSTL1. Both proteins are likely protease inhibitors, one for serine, one for thiol proteases. Both are likely secreted and seem to be overexpressed in testis tissue. Upon closer inspection the interface does not show good contacts. Nothing known about interactions between proteases. Not sure about this prediction.

http://prodata.swmed.edu/humanPPI/results/Q96BF3_Q9UM44 (S0_S0) 41 IFres
Interaction between TMIGD2 (Transmembrane and immunoglobulin domain-containing protein 2) and HHLA2 (HERV-H LTR-associating protein 2). This interaction seems well known and functionally described, is mentioned in Uniprot in TMIGD2 entry: "Through interaction with HHLA2, costimulates T-cells in the context of TCR-mediated activation." The predicted interface is between the extracellular Ig-like domain of TMIGD2 and the extracellular V-set/Ig-like domain of HHLA2. The interface consists of many contacts and is looking good upon closer inspection.

http://prodata.swmed.edu/humanPPI/results/Q6NTE8_Q96S96 (S0_S0) 40 IFres
Interaction between MRNIP (MRN complex-interacting protein) and PEBP4 (Phosphatidylethanolamine-binding protein 4). There is some evidence in STRING for these two proteins to interact with each other but I think it is a mistake. It is based on homology to yeast where there is a structure of the mitochondrial ribosome that is supposed to contain homologs of the two human proteins but I think this is a mistake for MRNIP. MRNIP functions in DNA repair by assisting MRN to execute its function in DSB repair. PEBP proteins are known to bind specific membrane lipids and therefore are known to function in cell signaling related to MAP or G protein of NF-KB or ERK. The interface also does not look very convincing.

http://prodata.swmed.edu/humanPPI/results/Q9UBM4_Q9UGK8 (S0_S0) 40 IFres
Interaction between OPTC (Opticin) and SERGEF (Secretion-regulating guanine nucleotide exchange factor). Opticin has some function in eye tissue preventing neovascularization, binds collagen fibrils. Is predicted to bind with its LRR-like repeat domain (LRRNT domain) to SERGEF and its WD-like propeller repeat domain. SERGEF is assumed to aid protein towards secretion. OPTC is a secreted protein. OPTC is also predicted to bind with this surface of its LRRNT domain to many other partner proteins with different folds. So, does not seem to be a very specific prediction. SERGEF has also quite some other modelled binding partners with different folds that are also predicted to bind to the same region on the WD repeat domain.

http://prodata.swmed.edu/humanPPI/results/P19022_Q8IYP2 (S1_S0) 37 IFres
Interaction between CDH2 (Cadherin-2) and PRSS58 (Serine protease 58). Cadherins are known to be cleaved to get fully functional. The N-terminal region is cleaved off increasing the adhesive function of extracellular CDH2. It homodimerizes and thereby mediates cell-cell contacts. But the cleavage is known to be done by furin-family proteases to which the serine proteases do not belong to. There are two predictions for both proteins. In one prediction the N-terminal domain of CHD2 is predicted to bind the protease and in the other prediction it is the C-terminal domain. Although the N and C-terminal domains of CDH2 are structurally related, they are predicted to bind to PRSS58 in different ways. It also doesn't look like that it predicts

how a substrate would bind to an enzyme. There is no other evidence for this interaction to exist. I think this prediction is very questionable.

http://prodata.swmed.edu/humanPPI/results/P07358_P10643 (S0_S2) 33 IFres
Interaction between C8B (complement component C8 beta chain) and C7 (complement component C7). This interaction is well known and also structurally studied. Both proteins are part of the MAC complex (membrane attack complex) which plays a role in innate immunity. There are a few structures solved, such as 8B0F that I looked at more specifically. There are two predictions for this interaction. One (S0_S1, so not this one selected here) corresponds to the resolved interface of both proteins in the MAC complex. The other prediction, so the one selected here, is a wrong prediction. The interface that C7 covers on C8B is covered by other subunits in the MAC complex. Protein C8B corresponds to chain D in 8B0F and C7 corresponds to chain C.

http://prodata.swmed.edu/humanPPI/results/P48304_Q16609 (S0_S0) 29 IFres
Interaction between REG1B (Lithostathine-1-beta) and LPAL2 (putative apoliprotein(a)-like protein 2). LPAL2 might be the product of a pseudogene. It has no evidence at protein level. REG1B is secreted and extracellular. The interface looks very poor upon closer inspection. This prediction is very questionable.

http://prodata.swmed.edu/humanPPI/results/Q03393_Q4G0W2 (S0_S0) 28 IFres
Interaction between PTS (6-pyruvoyl tetrahydrobiopterin synthase) and DUSP28 (dual specificity phosphatase 28). The structure prediction docks the C-terminal end with a Glu as last residue of PTS into the catalytic site of DUSP28. There is a structure resolved of DUSP28 that describes where the catalytic site is (5Y15). Other than that there are no good predicted residue-residue contacts between both proteins. Could be that the C-terminal Glu plus C-terminus are attracted in the structure prediction by the rather positively charged active site of DUSP28. Interestingly, the study that resolved the structure of DUSP28 mentions that the catalytic site is very narrow probably explaining the low enzymatic activity of DUSP28. Overall, I think this prediction is very questionable.

http://prodata.swmed.edu/humanPPI/results/O43609_O43610 (S0_S0) 26 IFres
Interaction between SPRY1 and SPRY3 (Protein sprouty homolog 1 and 3). Interaction is reported in STRING but just based on co-mentioning of homologs in abstracts. Interface is predicted between both SPROUTY domains which I think don't form stable folds on their own. They probably need the right binding partner. The structure that is predicted here though does not fit for stabilizing the folds because the interface is very very small. There is also not other evidence that these domains would homodimerize. No structures resolved for these proteins but quite some evidence that they bind other types of domains such as EVH1 domains. I think this prediction is likely wrong.

http://prodata.swmed.edu/humanPPI/results/Q8N6H7_Q9Y678 (S0_S1) 23 IFres
Interaction between ARFGAP2 (ADP-ribosylation factor GTPase-activating protein 2) and COPG1 (Coatomer subunit G1). This interaction has been experimentally described using

pulldown experiments and westernblotting. It is also functionally described that ARFGAP2 is the GAP for ARF1. Hydrolysis of the GTP bound to ARF1 by ARFGAP2 might regulate release of the coatomer complex from the Golgi. There are two predictions for this interaction. One involves a motif in ARFGAP2 that is predicted to bind via beta strand augmentation to a domain in COPG1. The sequence of the motif looks like a conserved island in the disordered region of ARFGAP2, so this looks like a true motif. However, this interface is not the one picked here, so I guess that this interface clustered with an interface from the PDB. Could we find out which one? The other prediction that is selected here involves the long helical repeat domain of COPG1 with various different disordered parts of ARFGAP2 but all predicted contacts are just a few, nothing that I think could mediate binding.

http://prodata.swmed.edu/humanPPI/results/P56537_Q9NWT1 (S0_S0) 21 IFres
Interaction between EIF6 (Translation initiation factor 6) and PAK1IP1 (p21-activated protein kinase-interacting protein 1). The prediction looks very interesting. It is a highly charged interface, primarily positive charges on EIF6 and negative charges on PAK1IP1 surface. EIF6 associates with the ribosome and regulates translation under stress conditions. Using structure 6LQM I found that the predicted interface with PAK1IP1 is compatible with binding of EIF6 to the ribosome. PAK1IP1 inhibits activation of the PAK1 kinase. There is some publication listed in Uniprot for the statement that PAK1IP1 might be involved in ribosome assembly, which is also regulated by EIF6 but I did not check this publication further. Based on uniprot there is no mention that PAK1 regulates ribosome activity. EIF6 is known to be regulated by phosphorylation but PAK1 is not mentioned in uniprot or when asking Perplexity. Both proteins are very small and the interface is predicted between two folded domains. According to STRING both proteins were found in a few studies in pulldown experiments but not clear if one of them was bait and the other prey.

Cluster DI49350_Humanppi_P54764_S1__Q15375_S2
Link for representative http://prodata.swmed.edu/humanPPI/results/P54753_P54764 does not work. It seems that in the more recent online version of the humanPPI resource this structure was gone but in the earlier downloaded version we have it is still there. This cluster contains 11 members without pLDDT filtering and 4 members with filtering. It consists of heterodimers of Ephrin-type A/B receptor family proteins. The proteins were fragmented in humanPPI and all predictions in this cluster contain the extracellular part (ectodomain). The predictions are made between the Ephrin LBD domain and the Ephrin receptor like domain on each chain. There is a structure resolved, 4BKF, which shows a very similar interface between a homodimer of EPHA4. Not sure why these interfaces were classified as novel? The predicted and resolved interface deviate a little in the contacts within the Ephrin receptor like domain starting at AA 203. Saved a chimera session file. It turns out that I looked at the wrong protein fragment combination S1-S1, and not S1-S2. The S1-S2 combination of protein fragments docks the intracellular kinase domain of one receptor against an extracellular fibronectin domain of the other chain, very unlikely to be true. I also added one example to the chimera session file.

Cluster DI37678_Humanppi_A0A1W2PR75_S0__Q9BQ65_S0

It contains 4 predicted interfaces between different SSU72-like proteins and USB1. SSU72 like proteins are phosphatases that dephosphorylate specific residues of the C-terminal domain (CTD) of RNAP2. USB1 is a phosphodiesterase or exonuclease that cleaves off adenines or uridines at the 3’ end of U6 snRNA to generate the mature and active form of these snRNAs. There are structures solved with USB1 binding to substrates (i.e. 6D31). SSU72 proteins are docked into the substrate binding pocket of USB1 with a surface that is shown in other resolved structures to be bound by Symplekin. This is another protein that is reported to be crucial for the activation of SSU72-like proteins and for directing them to RNAP2 CTD dephosphorylation. Because both surfaces, in USB1 and SSU72-like proteins are found to bind very different other surfaces I doubt that these predictions are true.

Cluster DI55236_Humanppi_Q8IZ81_S0__Q9NVJ2_S0
It is a cluster formed by interfaces between members of the ELMOD domain containing protein family and the ADP-ribosylation factor (ARF)-like protein family. There are functional links between both protein families in that ELMOD proteins, i.e. ELMOD1 are known to act as GAPs for the ARF family GTPases (https://doi.org/10.1074/jbc.M112.417477). I couldn’t find anything related in the PDB and am wondering which side of the interacting surfaces were found to be similar to an interacting surface in the PDB. Interactions between both proteins are important for ciliary function and mutations in these proteins are associated with human disease. The predicted interfaces look great. Lot’s of very plausible AA contacts. The surface in the PDB that it was similar to is from the structure 8IAH, chain X. This Tropomyosin, a filament protein consisting of one super long helix. It has absolutely no structural similarity to the ELMOD1 fold apart from one single helix in ELMOD1 that is at the interface. I guess this is the interface similarity that it found. So, this cluster is highly novel.

Cluster DI42185_Humanppi_O94973_S2__P61966_S0
This cluster contains interfaces between the alpha1/2 subunit of the AP-2 complex and the sigma subunit. There are structures resolved (i.e. 9PWA) that show how the sigma subunit closely binds to the N-terminal (S1 fragment) of the alpha subunit and this is correctly predicted by humanPPI. However, this putatively novel cluster of interfaces docks the sigma subunit against the C-terminal domain of the alpha subunit. Given the strong evidence for how the sigma subunit engages with the alpha subunit to form the AP-2 complex, I don’t think that this alternative predicted interface has biological relevance.

Cluster DI52161_Humanppi_Q14831_S1__Q14833_S2
This cluster contains interfaces between Metabotropic glutamate receptors. In humanPPI they fragmented the proteins based on two separate folds they have. Both folds have a small intramolecular contact as predicted by AlphaFold. Pairing of an N-terminal domain of one receptor with the C-terminal domain of another receptor docks both domains in the exact same way as seen in the monomer for both domains. There are multiple structures suggesting that these receptors heterodimerize via the N-terminal domain only. Therefore, I don’t think that these predictions are accurate.

Cluster DI37654_Humanppi_A0A1W2PQ27_S0__Q5H9L4_S0

This cluster contains interfaces between SSU72-like proteins 1/3 and TAF7L (Transcription initiation factor TFIID subunit 7-like). As mentioned already earlier, SSU72-like proteins form stable complexes with Symplekin. TAF7L is predicted to bind against that surface that is used by Symplekin. It only covers a smaller fraction of the surface on SSU72-like proteins and the predicted AA contacts do not look very convincing. I don't think that this is a good prediction. There are no structures available for TAF7L.

Cluster DI52067_Humanppi_Q14563_S0__Q9HCM2_S2
This cluster contains two interfaces between Plexin proteins and Semaphorin proteins. These protein families are known to interact with each other. A structure has been resolved of one example (6FKN). It seems that for the correct fragments (S0-S1) this interface is correctly predicted. However, in comparison to this, the interface predicted for fragments S0-S2 looks wrong.

Cluster DI53651_Humanppi_Q5W0V3_S0__Q9NUY8_S0
This cluster contains interfaces predicted between FHIP2A/B (FHF complex subunit HOOK interacting protein 2A) and TBC1D23 (TBC1 domain family member 23). Based on uniprot annotations it is hard to know if these proteins would do something functionally related but STRING reports a publication that mentions both in a proximity-based study (PMID:34882091), which might be worth a closer look. FHIP proteins function as part of a larger complex that seems to be involved in cargo transport along microtubules together with Dynein-1. A structure of this complex was recently resolved with FHIP1B (8QAT). Superimposing this structure with the models on the FHIP proteins one can observe that the other subunits of the complex (HOOK3, AKTIP) do not overlap in their interface with how TBC1D23 is predicted to bind to FHIP2A/B. The predicted interface is quite intensive with a lot of sensible AA contacts. I think this is a very interesting prediction.
We ran AF3 with FHIP1B, AKTIP, HOOK3, and TBC1D23 which resulted in a complex model where TBC1D23 is docked to a different surface on FHIP1B compared to how it was predicted to bind to FHIP2A/B. Closer inspection revealed that at the surface where TBC1D23 is predicted to bind FHIP2A/B, FHIP1B has a large, mostly disordered insertion, which might block that interaction surface or modify it. We ran AF3 again, this time with FHIP2B, AKTIP, HOOK3, and TBC1D23 and now, it docked TBC1D23 to the complex in the same way how it does it for FHIP2B and TBC1D23 alone. In the complex model, AF3 also predicts that the C-terminal fold of TBC1D23 binds on top of FHIP2B which is different to the humanPPI model where this domain does not bind anything. However, AF3 is not confident at all in this additional contact. The previously mentioned publication (PMID:34882091) explains more about what is known about the incorporation of the different FHIP proteins into the FHF complex and performed a systematic proteomics-based and functional study to figure out if the different FHIP proteins enable the FHF complex to recognize and transport different cargos, which based on their study, they do. They found TBC1D23 in their proximity proteomics data to only be close to FHIP2A/B but not FHIP1A/B. Along with other proximal proteins the data suggests that FHIP2A/B might more specifically function in endosome-Golgi transport compared to FHIP1A/B complexes.

Cluster DI56807_Humanppi_Q96LK0_S0__Q9UBK7_S0

This cluster contains interfaces predicted between the centrosomal protein CEP19 and the Rab-like proteins RABL2A/B. The interaction between both proteins is well studied according to uniprot and functions in ciliation. Both are small proteins with one fold although CEP19 has an IDR that already in the unbound state is predicted in a quite particular fixed conformation with some confidence. It is exactly the conformation in which it is predicted to wrap around the fold of RABL2A/B. I think this is a highly confidently predicted interface. Interestingly, looking up CEP19 in the AlphaFold database it presented to me also a predicted complex structure from NVIDIA with RABL2A (https://alphafold.ebi.ac.uk/entry/AF-0000000211785136).

DI49269_Humanppi_P54198_S1__Q9NPG3_S0
This cluster contains two interfaces between HIRA and UBN1/2 (Ubinuclein 1/2). Both proteins are known to be part of the same complex, the HIRA histone chaperone complex, formed in addition by CABIN1. The complex interacts with the histone chaperone ASF1A to mediate Histone H3.3 specific binding and chromatin deposition. There is a structure resolved between a small fragment of UBN1 in complex with H3, H4, and ASF1 (4ZBJ). The interfaces predicted here involve the N-terminal WD40 propeller domain of HIRA for which there is no structure resolved yet (however, as realized later, a structure exists of the yeast homolog HIR2). UBN1/2 is predicted to wrap around the WD40 domain binding at three occasions in beta strand augmentation to three propeller repeats. The AA contacts look quite convincing. In addition there is a larger loop predicted to bind to the top (or bottom) of the WD40 domain. However, this loop exactly corresponds to the short fragment of UBN1 that is resolved in the structure mentioned earlier (4ZBJ). In this structure this UBN1 fragment is resolved to bind in the exact same conformation as predicted here to H3/H4. I would therefore think that this part of the prediction is wrong, however, the beta strand augmentation interface on the side of the WD40 domain could be right. One should run an AF prediction of the whole HIRA complex together with H3, H4, and ASF1 to see which interfaces are then predicted. There is another structure resolved (5YJE) which resolved the C-terminal HIRA domain of HIRA and found that it trimerizes. The WD40 and HIRA domain are separated by an IDR. UBN1/2 is quite a disordered protein. The UBN1 fragment that binds H3/H4 is predicted by AlphaFold also in the monomeric state of UBN1 in this conformation. However, the region that is predicted to bind in beta strand augmentation has a different but still quite rigid conformation in the UBN1 monomer model. Inspecting the AF model of HIFA, UBN1, H3, H4, and ASF1A I can confirm that also in this larger complex AF predicts for that disordered bit of UBN1 to bind in beta strand augmentation to the propeller domain of HIRA, however, the top (or bottom) of the propeller domain is not interacting with UBN1 anymore but with H4. The interaction between ASF1A, H3, H4 and UBN1 is largely correctly predicted as determined upon superimposition with 4ZBJ. It is cool to see how the propeller domain is an integral part of the complex also predicted to interact with the fold of ASF1A. Unfortunately, AF3 also predicts huge alpha helical domains for the larger disordered regions of HIRA and UBN1. I colored them grey in the chimera session file. I think that the positioning of the C-terminal domain of HIR and of the folded region of UBN1 is probably wrong.
We had the idea to search with the probably correct HIRA-UBN1 interface (involving the beta-strand augmentation) and excluding the probably wrong part involving the top (or bottom) of the WD domain for similar interfaces in the PDB. This identified the structure 8GHN, a large

structure of the yeast HIRA complex. This structure also includes CABIN1 (human homolog of yeast HIR3). This structure shows the exact same beta strand augmentation of UBN1 binding to the WD domain of yeast HIR2.  The top (or bottom) surface of the WD domain of yeast HIR2 however interacts with HIR3 (human CABIN1) suggesting that the predicted contact with H4 is likely wrong. It is kind of crazy how AlphaFold is misguided by exposed interaction surfaces for which the correct partner is not provided in the input file. In conclusion, this interface is not novel and it was identified as novel because of the wrong secondary contact of UBN1 and the top (or bottom) of the WD domain of HIRA.

DI49733_Humanppi_P60842_S0__Q13347_S0
This cluster contains interfaces predicted between EIF4A1/2 and EIF3I, both are eukaryotic translation initiation factors, once from factor 3 and once from factor 4. Both subunits work in the scanning and recognition of the start codon in mRNAs. There is a structure resolved (8oz0) which has both factors. It shows though only a tiny contact between them. EIF4A1 is otherwise not connected at all to other subunits of the complex. I would think that some parts could not be properly assigned in the cryoEM, I am also not sure that EIF4A is accurately positioned? EIF3I only has one propeller domain. The predicted interface puts EIF4A on the opposite side of the propeller compared to what is seen in the structure. However, the predicted AA contacts are not very convincing. I am not sure that this is a correct prediction.
